# Leveraging genome-wide association studies and genomic prediction for distinctness, uniformity, and stability (DUS) testing in maize

**DOI:** 10.64898/2026.06.10.731330

**Authors:** Anurag Daware, Clemens Hacke, Márton Pécs, Arnaud Remay, Philipp Starnberger, Christina Schraml, Cécile Collonnier, François Laurens, Karl Schmid

**Affiliations:** University of Hohenheim, Stuttgart, Germany, Institute of Plant Breeding, Seed Science and Population Genetics; National Food Chain Safety Office (NEBIH), Hungary; Groupe d’Etude et de contrôle des Variétés Et des Semences (GEVES), France; Austrian Agency for Health and Food Safety (AGES); Community Plant Varieties Office (CPVO), Angers, France; French National Institute for Agriculture, Food, and Environment (INRAE); Cluster of Excellence GreenRobust, University of Hohenheim, Stuttgart, Germany

**Author notes:** Send correspondence to: Anurag Daware, & Karl Schmid,.

**Keywords:** Distinctness, uniformity, and stability testing, Plant variety protection, Genome-wide association study, Maize, Genomic prediction

## Abstract

Testing for distinctness, uniformity, and stability (DUS) is a requirement for plant variety registration and based on phenotypic traits, which is time-consuming and sensitive to environmental variation. Advances in genomics allow to complement DUS testing with molecular markers, for which two models in DUS testing were proposed by the Union for the Protection of New Varieties of Plants (UPOV). A use cases was described for maize, but an implementation has been hindered by a lack of suitable markers and validated analytical frameworks. We address these challenges by integrating historical DUS characteristics scores from 352 European hybrid maize varieties with high-density genome-wide single nucleotide polymorphism (SNP) data. Using genome-wide association studies (GWAS), we identified 18 genomic regions and candidate genes associated with 12 DUS characteristics, enabling the development of diagnostic markers consistent with the UPOV model “Characteristic-Specific Molecular Markers”. Since most DUS traits are polygenic, we combined GWAS-informed marker selection with XG-Boost-based machine learning to predict notes of DUS characteristics. This approach achieved strong predictive performance across multiple traits (mean accuracy 0.67), demonstrating its potential for managing reference collections under UPOV model “Combining phenotypic and molecular distances in the management of variety collections”. Both approaches were validated for two characteristics using independent public USDA-NPGS maize datasets (>1,700 accessions) highlighting the value of public data for method validation. We also identify key limitations of historical DUS data, including imbalanced and sparse trait representation, and discuss mitigation strategies. Despite these constraints, our results demonstrate that molecular markers may improve maize DUS testing, enabling faster, more accurate variety registration and supporting accelerated crop improvement.

**Key message:** Historical DUS datasets can be used to identify marker-trait associations of DUS characteristics using genome-wide association study (GWAS) and to develop a genomic prediction framework for an accurate prediction of DUS character notes from marker data.

## Introduction

Developing improved varieties is essential for enhancing agricultural productivity, ensuring food security, and addressing climate resilience challenges. Breeding new crop varieties requires substantial investments in time, financial resources and expert personnel (Shimelis and Laing 2012). To protect these investments and incentivize innovation, breeders rely on plant variety protection (PVP) systems that grant exclusive commercial rights (Yu and Chung 2021). The International Union for the Protection of New Varieties of Plants (UPOV) Convention establishes standardized criteria for variety protection through distinctness, uniformity, and stability (DUS) requirements (UPOV 2002). Despite its central role in PVP, traditional DUS testing which evaluates morphological characters of candidate varieties alongside a reference panel of registered varieties faces critical limitations (Yu and Chung 2021). The process is time-consuming, costly, and heavily reliant on expert judgment. Environmental factors including soil, climate, and seasonality can confound character expression, complicating efforts to distinguish genetic from environmental influences (Yang et al. 2021; Yu and Chung 2021). As breeding advances, many varieties exhibit increasingly subtle phenotypic differences that challenge visual or morphological distinction. Detecting essentially derived varieties (EDVs) with minor but legally significant differences presents additional difficulties. These constraints create an urgent need for more robust, efficient approaches to support DUS testing (Jamali et al. 2020).

Molecular markers like simple sequence repeats (SSRs) and single nucleotide polymorphisms (SNPs) have emerged as valuable tools to complement traditional DUS assessments (Gunjaca et al. 2008; Noli et al. 2008; Ibáñez et al. 2009; Jones et al. 2013). They offer high resolution, reproducibility, and environmental independence, enabling earlier and more objective evaluations. It was proposed that molecular markers are necessary to improve DUS testing efficiency (Mendler-Drienyovszki et al. 2024). The Biochemical and Molecular Techniques (BMT) Review Group of UPOV has approved two models for integrating molecular markers into DUS testing (UPOV (2020); CPVO (2017)). The first model “Characteristic-specific Molecular Markers” (thereafter Model 1) employs markers tightly linked to DUS characteristics as direct predictors trait notes. They are valuable when marker–phenotype associations are well-established and phenotypic scoring is challenging in the field. The second model “Combining phenotypic and molecular distances in the management of variety collections” (thereafter Model 2) calibrates genome-wide molecular differences against minimum phenotypic distances to establish molecular distinctness thresholds, helping manage reference collection size to reduce the size of field trials and the costs of DUS testing. This model expects a high correlation between molecular and phenotypic distances.

A successful implementation of both models requires high-quality marker selection, validation across diverse germplasm, standardized methodologies, and the establishment of molecular profile databases (Jamali et al. 2019; Yu and Chung 2021). Molecular marker-assisted DUS testing following accepted UPOV models has shown promising results across diverse crops including tomato, barley, and soybean (Guan et al. 2020; Zhang et al. 2020; Achard et al. 2020; UPOV 2021; Yang et al. 2021). Therefore, it is of great interest to evaluate this approach for major crops such as maize.

Europe harbors a remarkable maize diversity, with 6,232 varieties (predominantly hybrids) registered with the European Plant Variety Office (CPVO), granting EU-wide protection and marketing rights (European Commission 2025). This diversity spans varieties bred for silage, grain, and biomass production, displaying substantial variation in traits like grain type (dent versus flint), maturity, and plant pigmentation (Balconi et al. 2024). Since most European maize varieties are hybrids, molecular markers offer additional value by enabling registration and verification of parental inbred lines with novel combinations of DUS characteristics, as described in Example 1 for UPOV BMT Model 1 in Document TGP/15/3 (UPOV 2020). Despite these advantages, molecular markers are not widely used in maize DUS testing under both BMT models 1 and 2. Experiences from other crops such as barley, rice, tomato and wheat suggest significant value in using genome-wide association studies (GWAS) for identification of such markers, indicating their potential in maize (CPVO 2013, 2017; Jones and Mackay 2015; Yang et al. 2021; Liu et al. 2022; Zanella et al. 2026). An evaluation of UPOV BMT Model 2 in maize used small marker sets (24–384 markers) and limited variety collections did not reveal a strong correlation between morphological and genetic distance (Gunjaca et al. 2008; Guan et al. 2020). The French examination office GEVES genotyped 4,000 maize inbreds and 400 hybrids with 312 SNP markers. Similarly, no strong correlation between DUS characteristics and molecular markers, but a genetic distance threshold of 0.20 was proposed for excluding varieties from reference collections (UPOV 2014; Thomasset et al. 2015). Although these findings indicate a practical feasibility of BMT Model 2, they do not satisfy the core requirement i.e. a robust correlation between molecular and morphological distances. Larger marker panels or modified approaches like genomic prediction for DUS traits have successfully strengthened correlations between DUS characteristics and genetic distances in barley, rice and perennial ryegrass (Jones and Mackay 2015; Liu et al. 2022; NIAB 2025; Roberts et al. 2026), but such work is lacking in maize.

In the present study, we harnessed historical DUS records from 352 hybrid maize varieties provided by four European examination offices, and combined them with genome-wide molecular markers to dissect the genetic architecture of DUS characters and establish both a marker resource and an improved methodological framework for integrating molecular markers into maize DUS testing that may align with UPOV BMT models. This included the identification of marker-trait associations using GWAS for key DUS characters from the CPVO maize testing protocols as a foundation for developing diagnostic markers suitable for applications according to UPOV BMT Model 1 (CPVO 2001, 2010; UPOV 2020). Furthermore, we developed a two-step genomic prediction workflow that combines binary GWAS-based marker selection with XGBoost machine learning to predict DUS character notes from genome-wide SNP data, which achieved superior accuracy compared to earlier methods with implications for application of UPOV BMT Model 2 in maize.

## Materials and Methods

### Plant Material and DUS Data

As a part of the ‘Innovations in variety testing in Europe (INVITE)’ project, a panel of 352 European registered hybrid maize varieties was assembled by procuring seeds from four examination offices (EO): Austrian Agency for Health and Food Safety (AGES, Austria; 27 varieties), Groupe d’Etude et de contrôle des Variétés Et des Semences (GEVES, France; 104 varieties), Bundessortenamt (BSA, Germany; 45 varieties), and National Food Chain Safety Office (NEBIH, Hungary; 176 varieties) after obtaining breeders’ consent to genotype these varieties and anonymizing the identity of these varieties by examination offices. We further received historical notes of 40 DUS characteristics for these varieties from each EO. These notes were produced by the EOs during variety registration and obtained by transforming raw measurements and observations into standardized notes according to the maize DUS testing protocol (CPVO 2010). For this, numeric codes (notes) were assigned to qualitative characteristics (typically ranging 1–9), grouping quantitative measurements into defined ranges corresponding to specific note values, and scoring characters based on the observed degree or intensity of each characteristic. The characteristics included morphological, phenological, and physiological traits (Table 1) and were classified according to their mode of expression as quantitative, qualitative, or pseudo-qualitative, following UPOV terminology. Quantitative characteristics represent continuous variation measured or scored on a scale, qualitative characteristics comprise discrete states of expression, and pseudo-qualitative characteristics represent ordered categories assessed visually. For technical reasons, the amount of data obtained for each of the 40 characters differed among EOs: AGES provided final notes for 8 characters, GEVES for 39 characters, BSA for 20 characters, and NEBIH for 30 characters. After receiving the DUS data, the integration across EOs was performed by aligning character descriptors using standardized UPOV codes (CPVO 2010). Due to the variable amount of data provided by EOs there was significant variation in coverage of characteristics with a high proportion of >40% missing data for most characters across the final set of 352 varieties. We therefore decided to exclude DUS characters with ≤ 40% missing data from further analysis. The remaining characters were stratified into three subsets to balance data completeness and sample size: 1) High-confidence subset: 4 characters with <8% missing data (311 varieties), 2) Moderate-confidence subset: 5 characters with 8–19% missing data (233 varieties), 3) Low-confidence subset: 10 characters with 20–39% missing data (116 varieties). This stratification ensured robust sample sizes for further analyses and accounting in a transparent manner for data quality limitations.

**Table 1.** Description of maize DUS traits as per CPVO-TP/002/3 (2010) and their classification based on data availability and missingness. Expression types according to UPOV nomenclature: PQ - Pseudoquantitative, QN - Quantitative, QL - Quantitative. Traits are grouped by their proportion of missing data in to high-confidence (high-conf), moderate-confidence (mod-conf) and low-confidence (low-conf) groups for genetic analyses.

| UPOV No. | CPVO No. | Description | Expression type | Data available | Subset | Missing Data (%) |
| --- | --- | --- | --- | --- | --- | --- |
| 1 | 1 | First leaf: anthocyanin coloration of sheath | QN | No | NA | 69.14 |
| 2 | 2 | First leaf: shape of apex | PQ | No | NA | NA |
| 3 | 3 | Foliage: intensity of green colour | QN | No | NA | 90.50 |
| 5 | 4 | Leaf: angle between blade and stem | QN | yes | Mod-conf | 17.80 |
| 6 | 5 | Leaf: curvature of blade | QN | yes | Low-conf | 37.39 |
| 8 | 6 | Tassel: time of anthesis | QN | yes | High-conf | 5.93 |
| 9 | 7 | Tassel: anthocyanin coloration at base of glume | QN | yes | Mod-conf | 17.80 |
| 10 | 8 | Tassel: anthocyanin coloration of glumes excluding base | QN | yes | Mod-conf | 18.40 |
| 11 | 9 | Tassel: anthocyanin coloration of anthers | QN | No | NA | 51.34 |
| 12 | 10 | Tassel: angle between main axis and lateral branches | QN | yes | High-conf | 5.93 |
| 13 | 11 | Tassel: curvature of lateral branches | QN | yes | Mod-conf | 17.80 |
| 14 | 12 | Tassel: number of primary lateral branches | QN | yes | High-conf | 5.93 |
| 15 | 13 | Ear: time of silk emergence | QN | yes | High-conf | 6.53 |
| 16 | 14 | Ear: anthocyanin coloration of silks | QN | yes | Mod-conf | 11.87 |
| 17 | 15 | Stem: anthocyanin coloration of brace roots | QN | yes | Mod-conf | 18.40 |
| 18 | 16 | Tassel: density of spikelets | QN | No | NA | 74.78 |
| 19 | 17 | Leaf: anthocyanin coloration of sheath | QN | No | NA | 94.36 |
| 20 | 18 | Stem: antho-<br>cyanin col-<br>oration of in-<br>ternodes | QN | No | NA | NA |
| 21 | 19 | Tassel: length of<br>main axis above<br>lowest lateral<br>branch | QN | No | NA | NA |
| 22 | 20 | Tassel: length of<br>main axis above<br>highest lateral<br>branch | QN | yes | Mod-conf | 17.21 |
| 23 | 21 | Tassel: length of<br>lateral branch | QN | No | NA | 72.70 |
| 24.1 | 22.1 | Only inbred<br>lines and vari-<br>eties with ear<br>type of grain:<br>sweet or pop:<br>Plant: length | QN | No | NA | 78.04 |
| 24.2 | 22.2 | Only hybrids<br>and open-polli-<br>nated varieties,<br>excluding vari-<br>eties with ear<br>type of grain:<br>sweet or pop:<br>Plant: length | QN | yes | Low-conf | 29.08 |
| 25 | 23 | Plant: ratio<br>height of inser-<br>tion of peduncle<br>of upper ear to<br>plant length | QN | yes | Low-conf | 23.15 |
| 26 | 24 | Leaf: width of<br>blade | QN | No | NA | 71.22 |
| 27 | 25 | Peduncle:<br>length | QN | No | NA | 53.12 |
| 28 | 26 | Ear: length | QN | yes | Mod-conf | 17.21 |
| 29 | 27 | Ear: diameter<br>(in middle) | QN | yes | Low-conf | 17.21 |
| 30 | 28 | Ear: shape | QN | No | NA | 71.81 |
| 31 | 29 | Ear: number of<br>rows of grain | QN | yes | Mod-conf | 17.21 |
| 32 | 30 | Only vari-<br>eties with ear<br>type of grain:<br>sweet or waxy:<br>Ear: number<br>of colours of<br>grains | QL | No | NA | 78.04 |
| 33 | 31 | Only varieties<br>with ear type<br>of grain: sweet:<br>Grain: intensity<br>of yellow colour | QN | No | NA | NA |
| 34 | 32 | Only varieties<br>with ear type<br>of grain: sweet:<br>Grain: length | QN | No | NA | 78.04 |
| 35 | 33 | Only varieties<br>with ear type of grain: sweet:<br>Grain: width | QN | No | NA | NA |
| 36 | 34 | Ear: type of grain | PQ | yes | Mod-conf | 22.55 |
| 37 | 35 | Only varieties with ear type of grain: sweet:<br>Ear: shrinkage of top of grain | QN | No | NA | NA |
| 38 | 36 | Ear: colour of top of grain | PQ | yes | Low-conf | 22.55 |
| 39 | 37 | Excluding varieties with ear type of grain: sweet:<br>Ear: colour of dorsal side of grain | PQ | No | NA | NA |
| 40 | 38 | Only varieties with ear type of grain: pop: Type of popped grain | QN | No | NA | NA |
| 41 | 39 | Ear: antho-cyanin col-oration of glumes of cob | QN | No | NA | 40.06 |

### Genotyping

We genotyped one plant from each of the 352 maize hybrid varieties using two complementary approaches. Because seeds were available only as F1 hybrids and not as their parental components, all genotyped plants were expected to be highly heterozygous. Of the 352 varieties, 188 were genotyped using a 600K SNP array (Unterseer et al. 2014), while the remaining 164 varieties were genotyped by low-coverage whole-genome sequencing (lcWGS) at a mean coverage of 2x. For SNP array genotyping, seeds from 188 varieties were grown under greenhouse conditions for three weeks. Young leaf tissue was collected and sent to TraitGenetics, SGS Institut Fresenius GmbH (Gatersleben, Germany), for DNA extraction and genotyping using the Affymetrix Axiom 600K maize SNP array. Quality control and SNP calling were performed using Axiom Analysis Suite 2.0 (Affymetrix), following the Axiom Best Practices Workflow.

For lcWGS, genomic DNA was extracted from three-week-old leaf tissue of 164 varieties using the Blood Kit (A&A Biotechnology, Gdańsk, Poland). DNA quality and concentration were assessed with a NanoDrop spectrophotometer and a Qubit 2.0 fluorometer (Qubit dsDNA BR Assay Kit, Thermo Fisher Scientific, USA). Sequencing libraries were prepared in-house using a bead-linked transposome-based protocol (Jones et al. 2023). Library preparation steps including DNA normalization, tagmentation, barcoding, pooling, cleanup, and size selection were performed as per the published method and Supplemental Protocol v2.0 (Jones et al. 2023). Libraries were then sequenced on an Illumina NovaSeq 6000 platform using 150 bp paired-end sequencing chemistry. The raw sequencing reads were de-multiplexed and quality-filtered using fastp v1.0.1 with parameters ‘-q 30 -u 30 -l 100 –detect_adapter_for_pe -p -z 1. Filtered reads were subsequently aligned to the Mo17 reference genome (Zm-Mo17-REFERENCE-CAU-2.0) (Chen et al. 2023) using the BWA v0.7.17 aligner (Li and Durbin 2009). Variant calling was performed using the GATK v4.2.5 best practices workflow (McKenna et al. 2010; Grzybowski et al. 2023).

### Integration of genotyping data

To enable integration of the SNP array and lcWGS genotype datasets, we first converted SNP array variant coordinates from the B73 AGP_v2 reference genome to the Zm-Mo17-REFERENCE-CAU-2.0 assembly using a custom lift-over pipeline (Axiom-LiftOver-BLAST; Supplementary Information). To allow the use of SNP array data as a reference panel for imputing lcWGS genotypes, we further annotated array variants with genotype dosage and genotype probabilities using a custom Python script. These annotations are required for compatibility with imputation software that explicitly models genotype uncertainty in lcWGS data.

Genotype dosage, defined as the expected number of alternate alleles, was encoded as 0 for homozygous reference (0/0), 1 for heterozygous (0/1 or 1/0), and 2 for homozygous alternate (1/1). Genotype probabilities were assigned with complete certainty to reflect the high confidence of SNP array genotype calls: for dosage 0, Pr(0/0) = 1 and Pr(0/1) = Pr(1/1) = 0; for dosage 1, Pr(0/1) = 1 and Pr(0/0) = Pr(1/1) = 0; and for dosage 2, Pr(1/1) = 1 and Pr(0/0) = Pr(0/1) = 0. This approach provides a simplified representation of genotype uncertainty for hard-called SNP array genotypes, which lack native probabilistic measures.

For dataset integration, SNP array genotypes were first phased using SHAPEIT v4.0 (Delaneau et al. 2019) and subsequently imputed without an external reference panel using Beagle v4.0 (Browning and Browning 2007). The resulting imputed SNP array dataset was then used as a reference panel to impute lcWGS genotypes based on genotype likelihoods, also using Beagle v4.0. Finally, the imputed datasets were merged and filtered for variants with a minor allele frequency (MAF) > 0.05 using bcftools (Danecek et al. 2021) to generate the final genotype dataset.

### Public genotype and phenotype data (DUS)

Genotype data for the USDA National Plant Germplasm System (NPGS) collection (ZeaGBSv2.7) were obtained from the Maize Diversity Project website (https://panzea.org). Corresponding phenotypic data for three traits: kernel color, kernel type, and cob shape, were retrieved from the GRIN-Global database (https://npgsweb.ars-grin.gov/gringlobal). These traits correspond to CPVO DUS descriptors: Ear: color of top of grain (CPVO36), Ear: type of grain (CPVO34), and Ear: shape (CPVO28), respectively. After aligning and matching genotype and phenotype records, the final datasets comprised 1,746 accessions for kernel color, 1,778 accessions for kernel type, and 1,551 accessions for cob shape.

### Analysis of genetic relatedness

Principal component analysis (PCA) was performed with the INVITE genome-wide SNP dataset using PLINK v2.0 (Chang et al. 2015). The number of retained principal components (PCs) was determined by cumulative explained variance, retaining the minimum number accounting for ≤ 85% of total genetic variation. Discriminant analysis of principal components (DAPC) was then used to characterize genetic relatedness and assign individuals to predefined groups (examination offices) using the adegenet R package v2.1.11 (Jombart 2008). Model optimization was conducted by cross-validation, varying the number of retained PCs (n.pca) while fixing the number of discriminant functions (n.da) to the number of groups minus one. The optimal PC number was selected using the xvalDapc function based on 50 replicates with 90% of samples used for training and 10% for validation. Classification accuracy was further evaluated using leave-one-out cross-validation, in which each individual was sequentially excluded, the model refitted, and group membership predicted for the excluded sample. Overall and group-specific classification accuracies were calculated as the proportion of correctly assigned individuals. Posterior membership probabilities from the DAPC were visualized as admixture bar plots, representing the proportional assignment of each individual to the predefined groups.

### Genome-wide association study (GWAS)

DUS testing comprises a heterogeneous set of characteristics, including quantitative, qualitative, and pseudo-qualitative traits. Many of these characteristics are initially measured on continuous scales but are subsequently converted into discrete states of expression recorded as ordinal notes (scale 1–9) by examination offices (EOs) in accordance with CPVO technical protocols CPVO-TP/002/3 and CPVO-TP/002/2 (CPVO (2001)). The categorical nature of these data presents methodological challenges for genome-wide association studies (GWAS). To address these chal-lenges, GWAS was performed for DUS characteristics using three complementary analytical approaches.

First, GWAS was conducted using the Fixed and random model Circulating Probability Unification (FarmCPU) method implemented in the GAPIT R package v.3.5, which accounts for genetic structure and relatedness by incorporating kinship matrices and principal components (Liu et al. 2016; Wang and Zhang 2021). Second, to explicitly model the ordered structure of states of expression, ordered multinomial regression was applied using OrdinalGWAS.jl v0.7.2 (German et al. 2020). This approach exploits the ordinal scale of notes (1–9) and can increase power when allelic effects act consistently across successive states of expression. In both analyses, the first five principal components were included as covariates, and association results were visualized using CMplot (Yin et al. 2021).

Third, because not all DUS characteristics necessarily follow a monotonic ordinal structure, a systematic pairwise case–control GWAS approach was implemented using logistic regression in PLINK v2.0 (Chang et al. 2015). Each DUS characteristic was decomposed into binary contrasts between pairs of notes, allowing the identification of genetic effects specific to transitions between particular states of expression. This approach is particularly relevant for characteristics such as Ear: colour of top of grain (CPVO 36), which is determined by multiple independent biochemical pathways (Diepenbrock et al. 2021; Jiang et al. 2023). Due to limited sample sizes within individual binary contrasts, these analyses were not used for candidate gene identification but instead served as input features (Cutoff value: *p* < 1 × 10^−5^) for downstream machine-learning-based genomic prediction models.

Candidate gene identification was based on FarmCPU and OrdinalGWAS results. For the latter methods, quantitative trait loci (QTL) were delineated using local score analysis applied to the corresponding *p*-value distributions following Fariello et al. (2017), implemented in R with *ξ* = 1 and Gumbel thresholds at *⍺* = 0.05 and 0.01. QTL were defined as contiguous chromosomal regions where the Lindley process exceeded the significance threshold, and these intervals were subsequently used for candidate gene identification.

### Genomic prediction of DUS characteristics

In accordance with UPOV BMT Model 2, which permits the use of molecular markers to support the management of reference collections by predicting the expression of DUS characteristics, we implemented a two-stage genomic prediction framework integrating GWAS-based marker selection with machine-learning classification (Figure 1). This approach aims to predict states of expression (notes) of DUS characteristics from genome-wide marker data, thereby assisting in the efficient organization and comparison of candidate varieties within reference collections.

**Figure 1.**
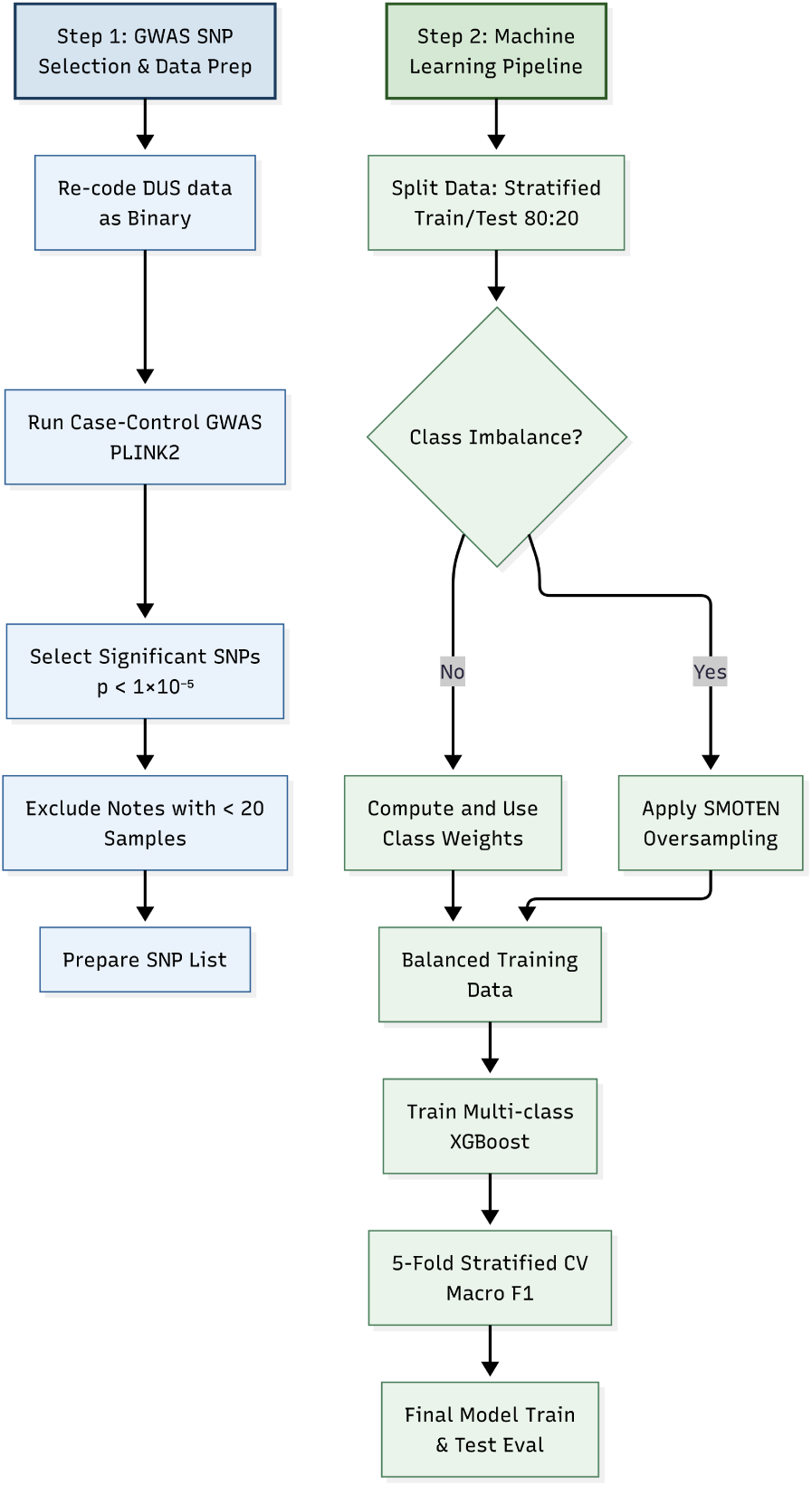
Schematic representation of the two-step genomic prediction workflow implemented for the genomic prediction of DUS characters.

As described above, pairwise case–control GWAS was conducted by re-coding each ordinal DUS characteristic into binary contrasts between two states of expression (notes) at a time and applying logistic regression in PLINK v2.0 (Chang et al. 2015). For each characteristic, SNPs showing significant association in any contrast (Cutoff value: *p* < 1 × 10^−5^) were extracted from the full genotype dataset to form reduced marker sets for prediction. This liberal threshold was chosen to capture informative molecular variation relevant to DUS characteristics while accounting for the multiple testing inherent to decomposing categorical characteristics into multiple binary contrasts. For model training, notes represented by fewer than 20 varieties were excluded to ensure reliable estimation. The remaining dataset was split into stratified training and test sets (80:20), preserving the proportional representation of states of expression. Class imbalance, which is common in DUS datasets derived from historical examination records, was addressed by applying class weights during model fitting. In cases of severe imbalance, the Synthetic Minority Oversampling Technique for nominal features (SMOTEN) was used to augment underrepresented states of expression (Chawla et al. 2002).

Multi-class prediction models were trained using XGBoost with SNP markers as predictors, using 200 trees, a maximum tree depth of 5, a learning rate of 0.1, 80% subsampling of individuals, and 80% subsampling of markers per tree (Chen and Guestrin 2016). Model performance was evaluated using stratified five-fold cross-validation, with the macro F1-score used as the primary metric, reflecting balanced performance across states of expression. Final models were trained on the full training set and evaluated on the independent test set, reporting overall accuracy, balanced accuracy, macro- and weighted F1-scores, and confusion matrices for each DUS characteristic. For each characteristic, multi-class receiver operating characteristic and precision–recall curves were generated to further assess predictive performance across states of expression.

## Results

### Data curation and DUS characteristic selection from historical DUS testing data

Historical DUS data were compiled for 352 maize varieties, comprising 33 DUS characteristics evaluated by four European examination offices (EOs). The integration of historical DUS data posed several challenges related to data harmonization and suitability for genetic analyses. A major limitation was the lack of original raw measurements, which are routinely converted by EOs into discrete notes (typically on a 1–9 scale) using office-specific grading procedures (e.g. standard deviation, least significant difference, or multiple comparison tests). As only final notes were available, differences in environmental conditions and assessment procedures could not be accounted for, and standardization across testing sites and years was not possible. The conversion of continuous measurements into discrete notes further reduced phenotypic resolution, potentially masking subtle differences among varieties. Moreover, the absence of replication across years or locations resulted in a single observation per DUS characteristic and variety, preventing estimation of heritability or correction for environmental effects. Nevertheless, because these scores are derived from standardized plant variety testing protocols, they provide robust and legally recognized descriptors of phenotypic characteristics.

Additional challenges arose from differences in CPVO technical protocols applied by the EOs. While AGES, BSA, and NEBIH followed the updated protocol CPVO-TP/002/3 (2010), GEVES applied the earlier version CPVO-TP/002/2 (2001). As a consequence, characteristics had to be aligned based on their descriptions, and incomplete overlap between protocols resulted in substantial missing data. Across DUS characteristics, the proportion of missing data ranged from 5% to 78% (Figure 2, Table 1).

**Figure 2.**
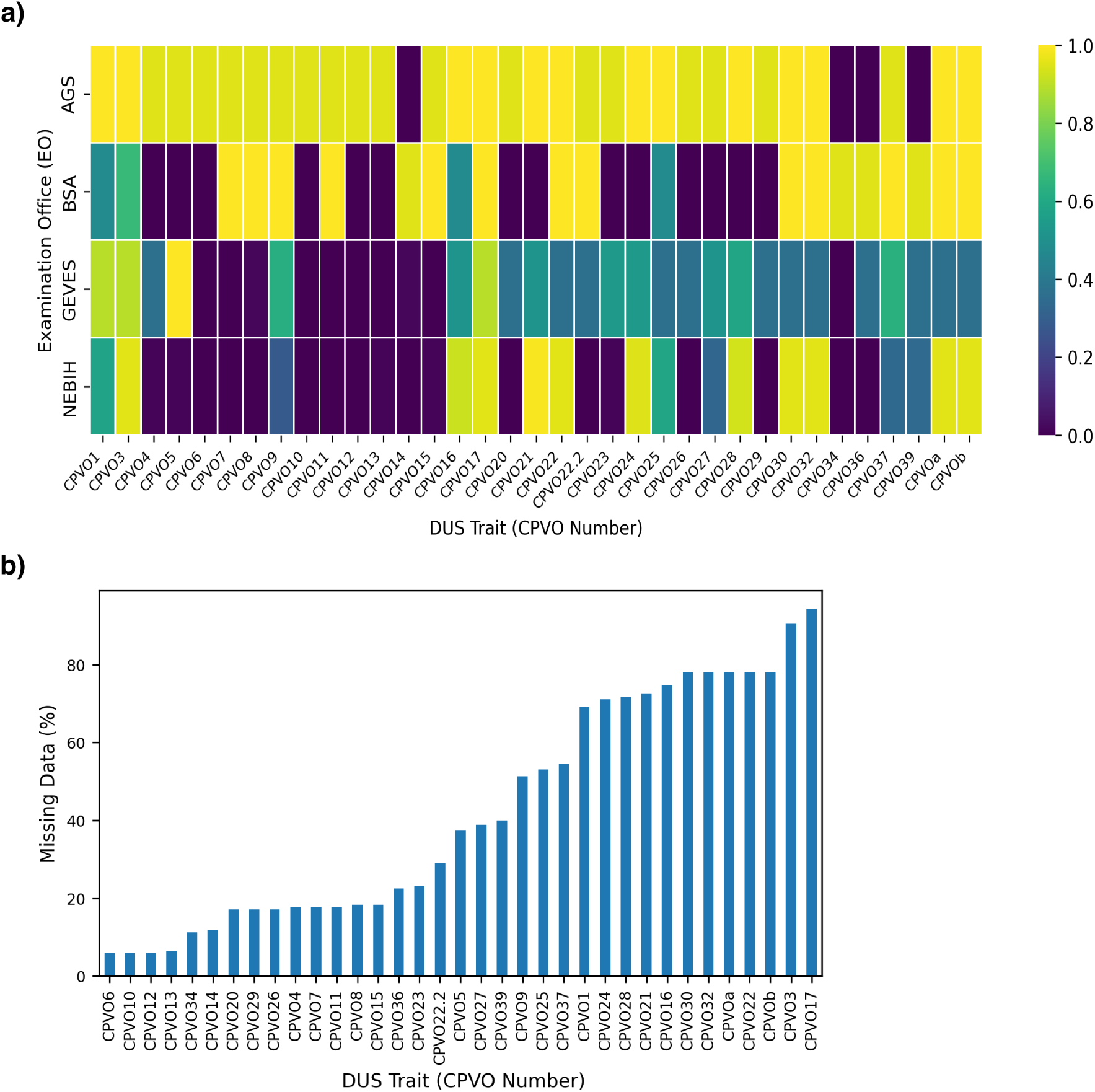
(a) Proportion of missing data for each DUS characteristic (CPVO number) across examination offices. Each cell shows the average proportion of missing data for a given combination of characteristic and examination office. (b) Percentage of missing data for each DUS characteristic (CPVO number) across all varieties, ordered from lowest to highest.

Finally, most DUS characteristics showed highly unbalanced distributions of varieties across notes, and for several characteristics certain states of expression were entirely absent (Figure 3). For example, the characteristic “Ear: colour of top of grain (CPVO 36)” comprises ten notes according to CPVO-TP/002/3, yet only four notes were represented in the present dataset, with two notes accounting for the majority of varieties. Similarly, for the anthocyanin-related characteristics “Tassel: anthocyanin coloration at base of glume (CPVO 7)” and “Tassel: anthocyanin coloration of glumes excluding base (CPVO 8)”, most varieties were assigned to the lowest state of expression, with few observations in higher categories. Such imbalanced note distributions reduce statistical power for detecting associations with less frequent states of expression and preclude the identification of marker sets capable of discriminating all notes of a characteristic. Consequently, these limitations restrict the suitability of historical DUS data for the development of diagnostic markers under UPOV BMT Model 1 that discriminate between all score categories, although they remain informative for exploratory analyses and predictive applications under Model 2.

**Figure 3.**
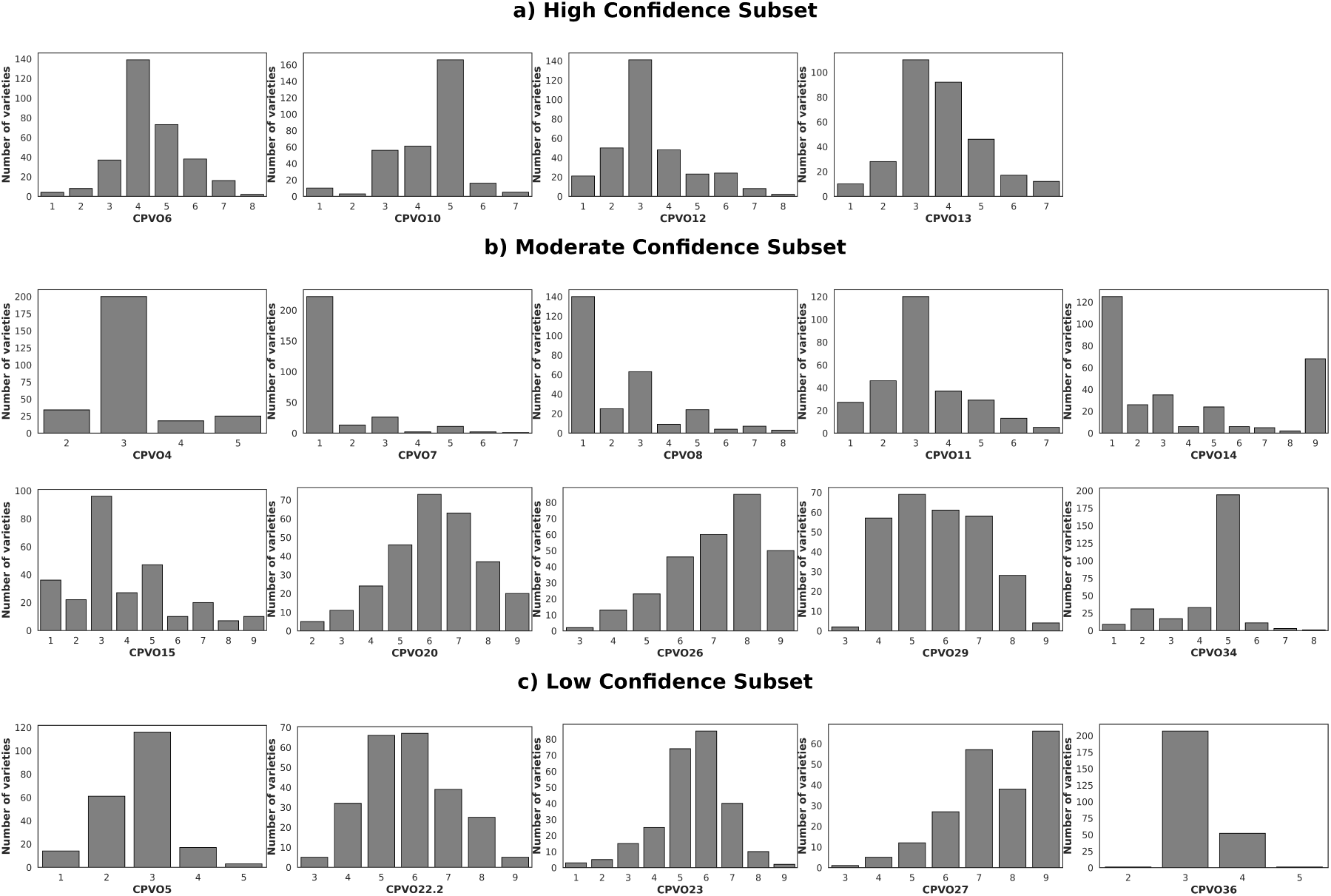
Distribution of notes across 19 DUS characters classified in three subsets based on level of missing data. High-confidence subset: a) 4 characters with <8% missing data across 311 varieties; b)Moderate-confidence subset: 5 characters with 8–19% missing data across 233 varieties; c) Low-confidence subset: 10 characters with 20–39% missing data across 116 varieties.

### Genetic variation and genetic relatedness in European maize varieties

To characterize genome-wide genetic variation among European maize varieties, 352 varieties examined by four examination offices (EOs) (AGES, BSA, GEVES, and NEBIH) were genotyped using a combination of SNP array genotyping and low-coverage whole-genome sequencing (lcWGS), followed by genotype imputation to integrate both datasets. This approach resulted in a final dataset comprising 466,363 high-quality SNPs distributed across the genome. Imputation was performed using SNP array–genotyped varieties as a reference panel and was based on genotype likelihoods to account for uncertainty associated with low sequencing coverage (see Materials and Methods). Because all varieties were F1 hybrids, heterozygosity levels were compared between lcWGS-imputed and SNP array–genotyped samples to assess genotyping and imputation consistency. Mean heterozygosity was slightly lower in lcWGS-imputed varieties (32.98%) than in SNP array–genotyped varieties (36.28%; Wilcoxon test: W = 7104, *p* < 2.2 × 10^−16^), likely reflecting challenges in detecting heterozygous loci in low coverage sequencing data and increased imputation error rates (Supplementary Figure 1) (Kardos and Waples 2025).

To assess the relationship between genetic structure and EO designation, principal component analysis (PCA) followed by discriminant analysis of principal components (DAPC) was performed (Figure 4). PCA revealed four partially overlapping genetic clusters that did not correspond directly to EO assignments (Figure 4 a). A subsequent DAPC analysis, based on the first five principal components explaining 85% of the total genetic variance, increased separation among EO groups (Figure 4 b). Varieties examined by NEBIH formed a compact and well-separated genetic cluster, characterized by minimal overlap with other EO groups and the smallest 95% confidence ellipse, indicating a high degree of genetic homogeneity. This distinctiveness was reflected in the highest leave-one-out cross-validation accuracy (93.1%), demonstrating that varieties tested by NEBIH could be reliably distinguished based on genome-wide marker data. In contrast, varieties examined by AGES, BSA, and GEVES occupied overlapping regions of multivariate genetic space, with broader confidence ellipses and substantial inter-group overlap, suggesting a shared genetic background or related breeding programs. Correspondingly, cross-validation accuracies were moderate for GEVES (62.0%) and BSA (64.0%), whereas AGES varieties showed lowest classifications accuracy (51.9 %), indicating a lack of EO-specific genetic separation. Posterior membership probabilities from the DAPC analysis further supported these patterns (Figure 4 c). NEBIH-tested varieties showed consistently high assignment probabilities (>75%) to the NEBIH cluster, with little evidence of admixture, whereas AGES, BSA, and GEVES varieties exhibited pronounced admixture, with individual varieties displaying substantial overlap with clusters of other EOs.

**Figure 4.**
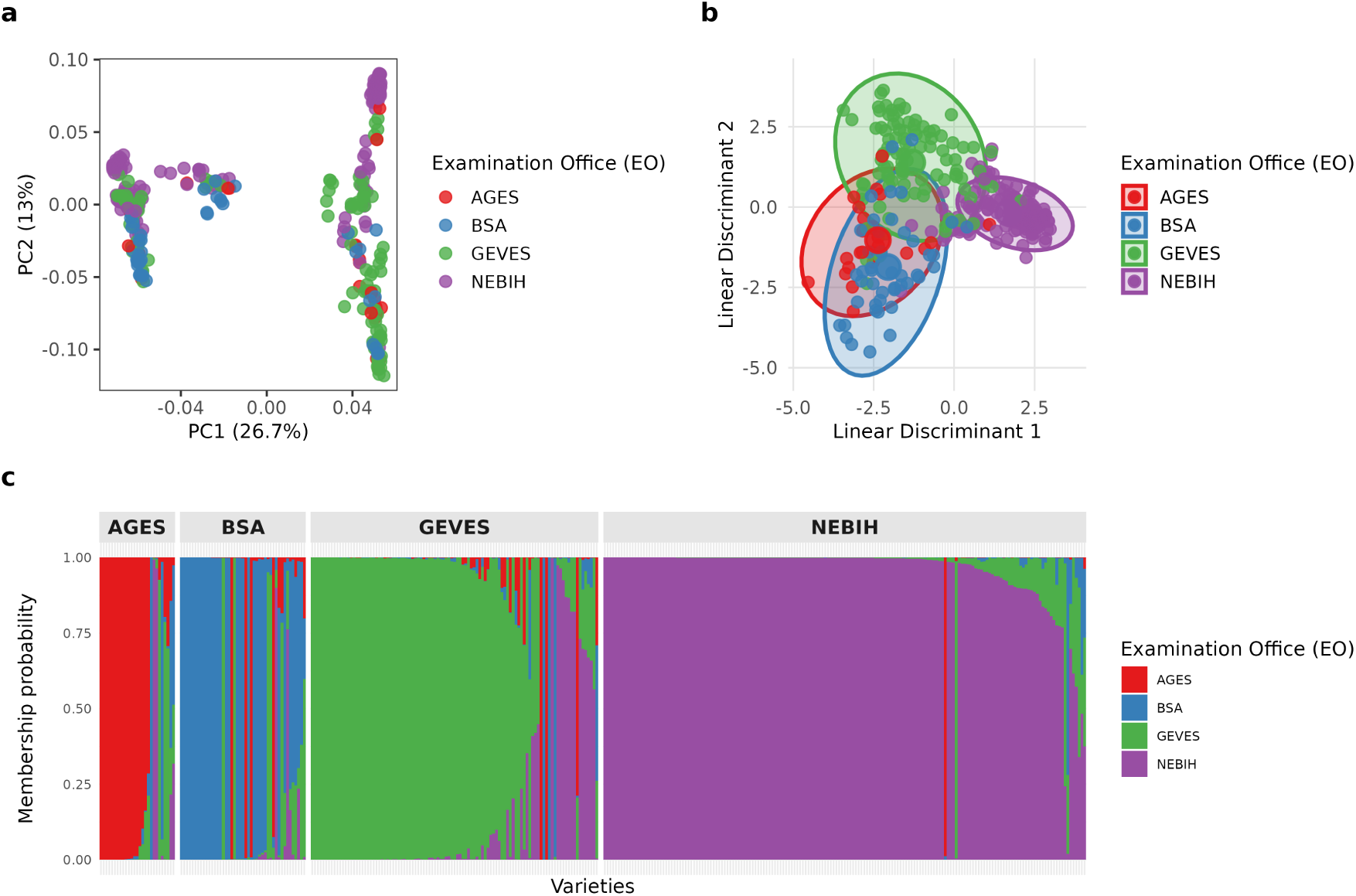
Genetic relatedness analysis of European maize (INVITE) varieties: (a) Principal component analysis (PCA) illustrating the broad genetic relatedness of European maize varieties. The groups correspond to four European examination offices (EOs) : AGES (Austria; *n* = 27), BSA (Germany; *n* = 45), GEVES (France; *n* = 103), and NEBIH (Hungary; *n* = 173). (b) Discriminant analysis of principal components (DAPC) using four European examination offices (EOs) as groups. (c) Admixture plot based on DAPC posterior membership probabilities showing genetic composition of each individual as a barplot of probabilities for assignment to four examination offices.

### Genetic architecture of DUS characters in maize

To identify marker–trait associations relevant for potential application under UPOV BMT Model 1 and to account for the ordinal structure and possible category-specific genetic effects of DUS characteristics, genome-wide association analyses were performed using three complementary approaches: (i) the Fixed and random model Circulating Probability Unification (FarmCPU) method, which accounts for population structure and kinship; (ii) OrdinalGWAS, which explicitly models the ordered nature of states of expression (notes); and (iii) binary logistic regression of pairwise contrasts between notes to examine specific transitions between states of expression (see Materials and Methods).

Candidate gene identification focused primarily on results from FarmCPU and OrdinalGWAS. Across all three confidence subsets, these analyses identified a total of 612 significant SNP associations (family-wise error rate of 5%) for 17 of the 19 DUS characteristics analyzed, including 73 associations detected by FarmCPU, 532 by OrdinalGWAS, and seven shared between both models (Supplementary Table 1). OrdinalGWAS detected a substantially larger number of associations, consistent with its design for ordinal phenotypes (German et al. 2020). Differences in the number of detected associations likely reflect differences in how the two models account for genetic relatedness. While OrdinalGWAS includes principal components as fixed effects, it does not explicitly model kinship, which may result in residual confounding. This was supported by quantile–quantile plots showing greater *p*-value inflation for OrdinalGWAS than for FarmCPU, suggesting more conservative control of confounding in the FarmCPU framework (Figure 5, Supplementary Figures 3 - 5). Consequently, OrdinalGWAS results were interpreted cautiously.

**Figure 5.**
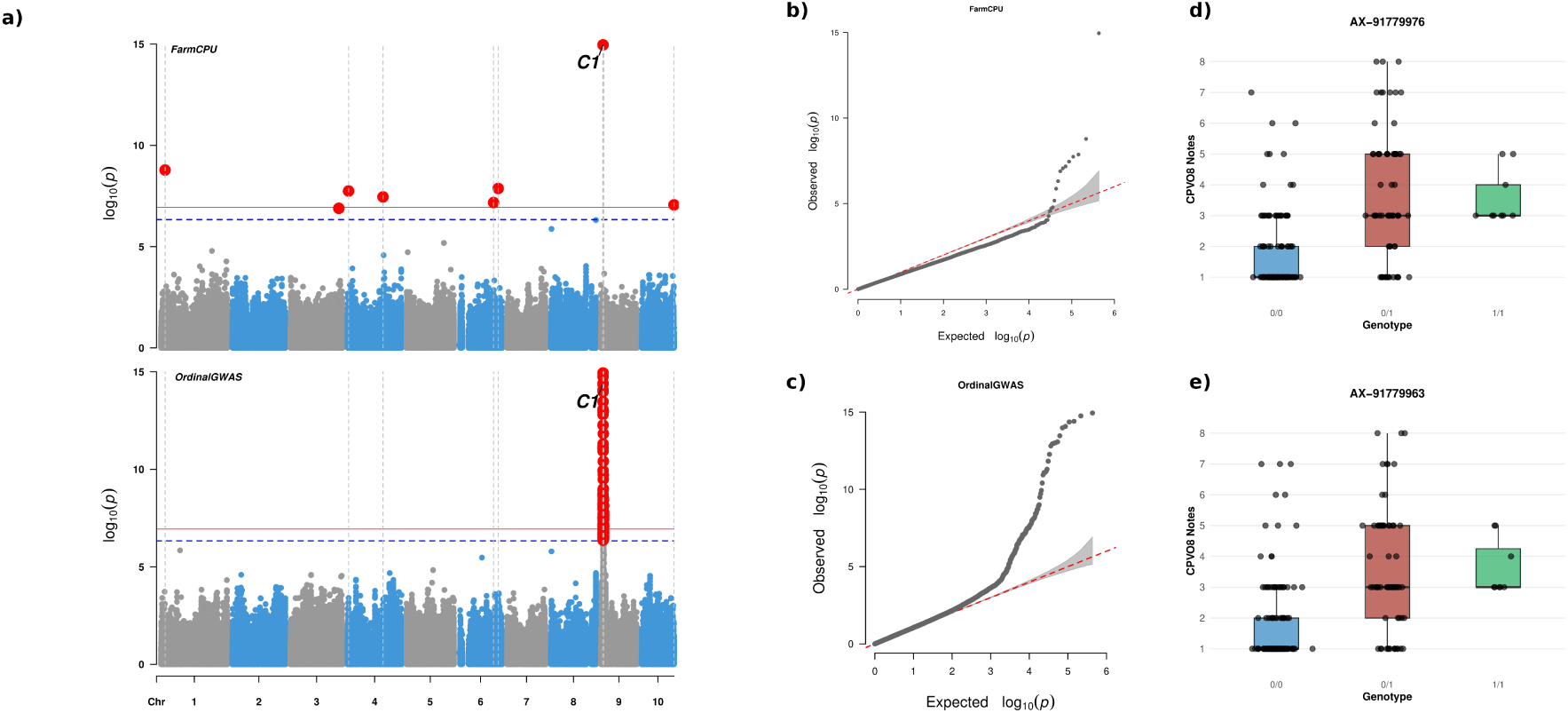
Genome-wide association analysis of the DUS characteristic “Tassel: anthocyanin coloration of glumes excluding base (CPVO 8)” in the INVITE maize dataset (*N* = 237). (a) Manhattan plots showing genome-wide SNP positions and association signals expressed as −log_10_(*p*) values from the FarmCPU (top) and OrdinalGWAS (bottom) analyses. The red solid line indicates the Bonferroni-corrected genome-wide significance threshold (family-wise error rate, FWER = 5%; *p* = 1.16 × 10^−7^), and the blue dashed line indicates a suggestive threshold (FWER = 20%; *p* = 4.62 × 10^−4^). Vertical dashed lines mark SNPs that were significant at FWER = 5% in at least one of the two models. (b,c) Quantile–quantile (QQ) plots comparing observed and expected *p*-value distributions for FarmCPU (top) and OrdinalGWAS (bottom). (d,e) Boxplots showing the distribution of states of expression (notes) for CPVO 8 across the three genotypic classes at the most significant SNPs identified by FarmCPU (top) and OrdinalGWAS (bottom).

After linkage disequilibrium–based pruning of associated SNPs (<10 kb), 351 distinct loci were retained, comprising 73 loci identified by FarmCPU, 271 by OrdinalGWAS, and seven shared between both models (Supplementary Table 2; Supplementary Figure 2). Both approaches identified multiple loci per DUS characteristic, with up to 10 loci detected for a single characteristic using FarmCPU (CPVO 34) and up to 63 loci using OrdinalGWAS (CPVO 6). These results point to a predominantly polygenic architecture; however, it is important to consider that environmental and protocol-related heterogeneity in DUS data may have inflated the number of detected loci (Supplementary Figures 3 - 5). Genomic regions with significant marker-trait associations were classified into those located near genes with known functions related to the corresponding DUS characteristics and those without obvious candidate genes (Supplementary Table 2). While tightly linked SNPs may be informative as predictive markers even in the absence of functional annotation, none of the identified variants showed sufficient allele specificity to reliably discriminate among all notes of a given DUS characteristic (Figure 5). Accordingly, subsequent analyses focused on well-characterized candidate genes as targets for the development of character-specific markers.

The value of a gene-focused interpretation of GWAS results is illustrated by a strong association detected for the DUS characteristic “Tassel: anthocyanin coloration of glumes excluding base (CPVO 8)” on chromosome 9. Both FarmCPU and OrdinalGWAS identified the same genomic region, which contains the well-characterized *colourless1* (*C1*) gene (Zm00014ba395290). In both analyses, the two most significant SNPs (lead SNPs; chr 9:10,739,095 and chr9:10,738,742) were located approximately 15.7 kb and 16 kb downstream of *C1 respectively* (Figure 5). The *C1* gene encodes a MYB transcription factor that regulates the expression of genes encoding key enzymes of the anthocyanin biosynthesis pathway (*A1*, *A2*, *Bz1*, *Bz2*, *Pr1*) and is known to control tissue-specific pigmentation in maize (Cone et al. 1986; Sainz et al. 1997).

Given the rapid genome-wide linkage disequilibrium (LD) decay in the GWAS panel (approximately 10 kb), the physical distance between the lead SNPs and the *C1* gene complicates candidate gene assignment based solely on proximity. The local LD patterns around the lead SNP (Chr9:10,739,095) revealed a 27 kb haplotype block. Notably, this block contains the C1 gene, suggesting it may be the functional driver of the observed association (Figure 6) (Gabriel et al. 2002; Han et al. 2018).

**Figure 6.**
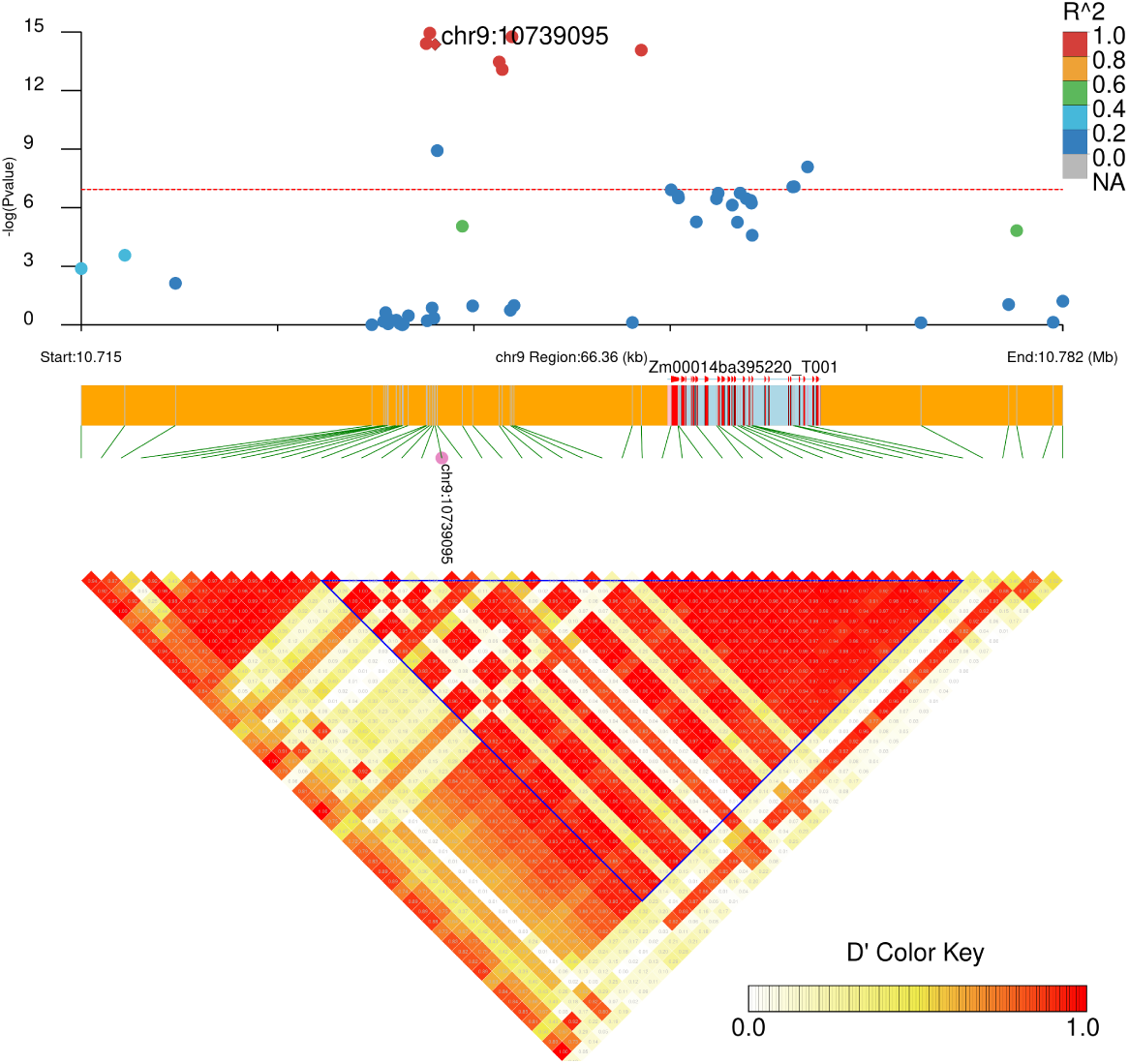
Linkage disequilibrium (LD) structure of genomic interval on chromosome 9 with the identified QTL for DUS character “tassel: anthocyanin coloration of glumes excluding base (CPVO8)”. The Manhattan plot shows a genomic SNP positions versus *p*-values expressed as – *log*_10_(*p*) from OrdinalGWAS, the red dotted line indicate the Bonferroni significance threshold (FWER = 5%; *p* = 1.16 × 10^−7^). The genomic region surrounding the lead-SNP (chr9:10739095) is visualized with its local LD and haplotype block structure. The lead-SNP is highlighted by a diamond-shaped marker in the upper association track. All neighboring SNPs are color-coded based on their pairwise linkage disequilibrium *r*^2^ with the lead-SNP, indicating the strength of their statistical correlation. The lower panel displays an LD heatmap calculated using *D*^′^, illustrating the historical recombination patterns and local LD architecture. The haplotype block containing the lead-SNP is delineated by a blue-bordered grid.

To further validate the association between lead-SNP and *C1* gene, we applied the local score approach, which aggregates association signals across adjacent markers by converting single-marker *p*-values into scores and identifying genomic regions with elevated cumulative evidence using a Lindley process (Fariello et al. 2017; Bonhomme et al. 2019). Using OrdinalGWAS-derived *p*-values and a tuning parameter of *ξ* = 1, the local score analysis identified a QTL on chromosome 9 spanning 4.7 Mb (Chr 9:10,349,281–15,123,800), which includes the *C1* gene (Figure 7 Supplementary Table 3). This result supports *C1* as the most plausible candidate gene underlying variation in this DUS characteristic and demonstrates the utility of the local score approach for delineating QTL intervals in panels with rapid LD decay. In contrast, no QTL was detected using FarmCPU-derived *p*-values, which is expected given the iterative conditioning of markers in FarmCPU that disrupts the continuous LD signal required for local score detection (Liu et al. 2016; Wang and Zhang 2021).

**Figure 7.**
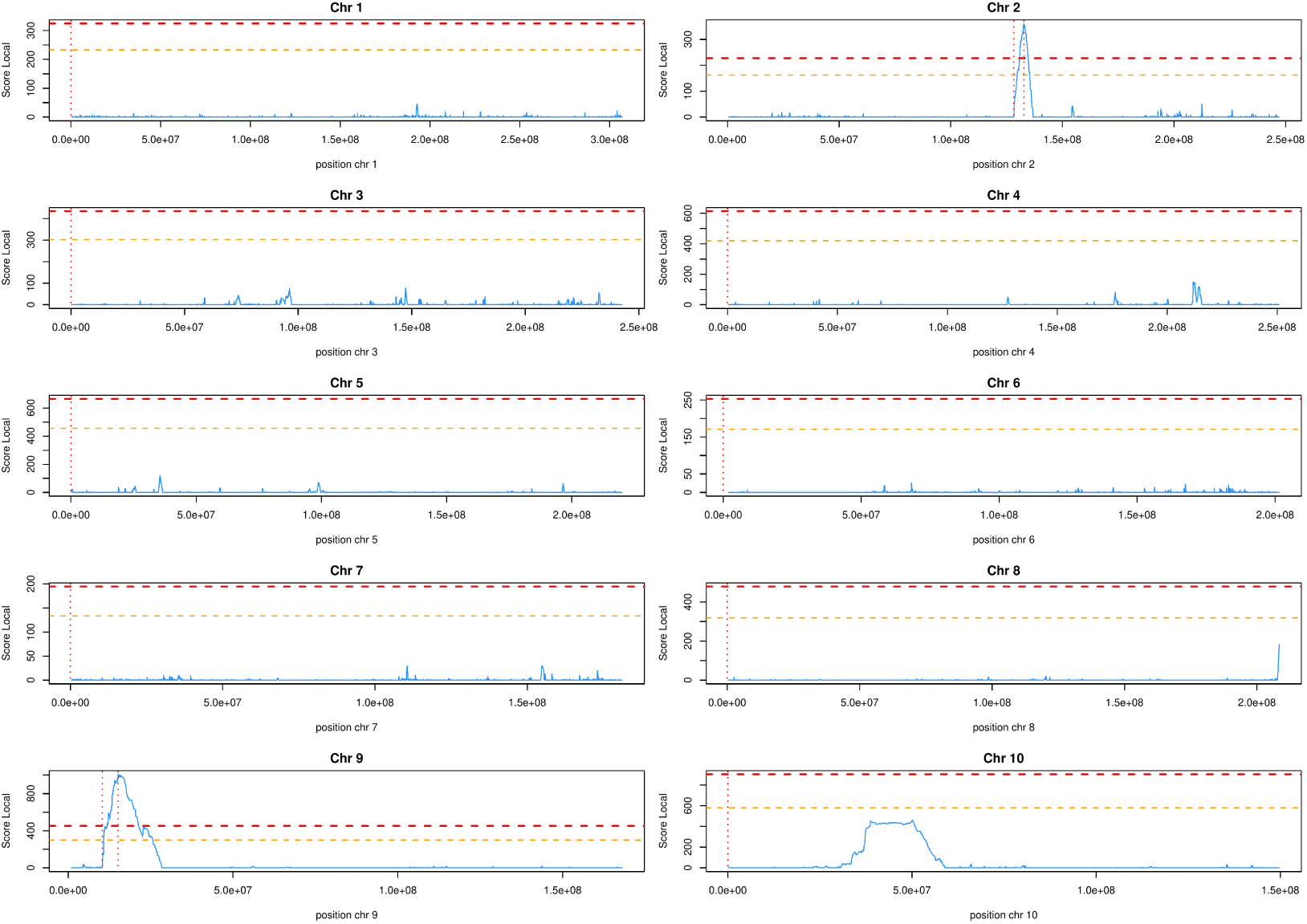
Detection of quantitative trait loci (QTL) associated with the DUS characteristic “Tassel: anthocyanin coloration of glumes excluding base (CPVO 8)” using the local score approach. The line plot shows values of the Lindley process (local score) computed from OrdinalGWAS *p*-values using a shape parameter of *ξ* = 1. Horizontal dashed lines indicate genome-wide significance thresholds derived from the Gumbel extreme value distribution at *⍺* = 0.05 (orange) and *⍺* = 0.01 (red). Vertical red lines delineate the boundaries of the detected QTL region.

Comparison of LD structure (*D*^′^) with local score profiles further illustrated the advantages of the local score approach. Although the 4.7 Mb region showed consistent association signals, haplotype blocks defined by *D*^′^ failed to capture this extended signal, whereas the local score remained elevated across the entire interval, more accurately reflecting the cumulative evidence for association (Figure 6).

### Analysis of public USDA data

To evaluate whether increased sample size and more balanced representation of states of expression (notes) improve the detection of genetic determinants of DUS characteristics, we analyzed publicly available data from the USDA–NPGS maize collection. Three characteristics routinely used in DUS examination were considered: “Ear: type of grain” (similar to CPVO 34), “Ear: shape” (similar to CPVO 28), and “Ear: colour of top of grain” (similar to CPVO 36). Due to differences in scoring protocols, this dataset was analyzed independently of the INVITE dataset.

The USDA–NPGS dataset provided substantially greater statistical power due to its larger sample size, although representation of states of expression remains unbalanced (Figure 8). Genome-wide association analyses identified 391 significant SNPs across the three characteristics, which were reduced to 250 independent genomic regions after linkage disequilibrium–based pruning (<10 kb). For the trait ‘kernel colour’, which aligns aligned with CPVO 36, several strong association signals co-localized with known genes of the carotenoid biosynthesis pathway (Figure 9, Figure 10, Supplementary Table 4 and 6). A major locus on chromosome 6 encompassed *YELLOW ENDOSPERM 1* (*Y1 : Zm00001d036345*), with the lead SNP located within the fourth exon. *Y1* encodes phytoene synthase, a key enzyme in carotenoid biosynthesis that determines yellow endosperm pigmentation (Buckner et al. 1990). On chromosome 2, the SNP with the highest *p*-value maps to second exon of *ZEP1* (*Zm00001d003512*), which encodes ZEAXANTHIN EPOXIDASE 1. This enzyme converts zeaxanthin to violaxanthin, modulating carotenoid composition and contributing to the orange-to-yellow coloration spectrum (Owens et al. 2019; Diepenbrock et al. 2021; Chen et al. 2024; Amoah et al. 2025). A third locus on chromosome 7 encompasses *DXS2* (*Zm00001d019060*), which encodes 1-deoxy-D-xylulose 5-phosphate synthase 2, the enzyme catalyzing the rate-limiting first step of the methylerythritol phosphate (MEP) pathway that supplies essential precursors for carotenoid synthesis (Cordoba et al. 2011; Fang et al. 2020; Diepenbrock et al. 2021).

**Figure 8.**
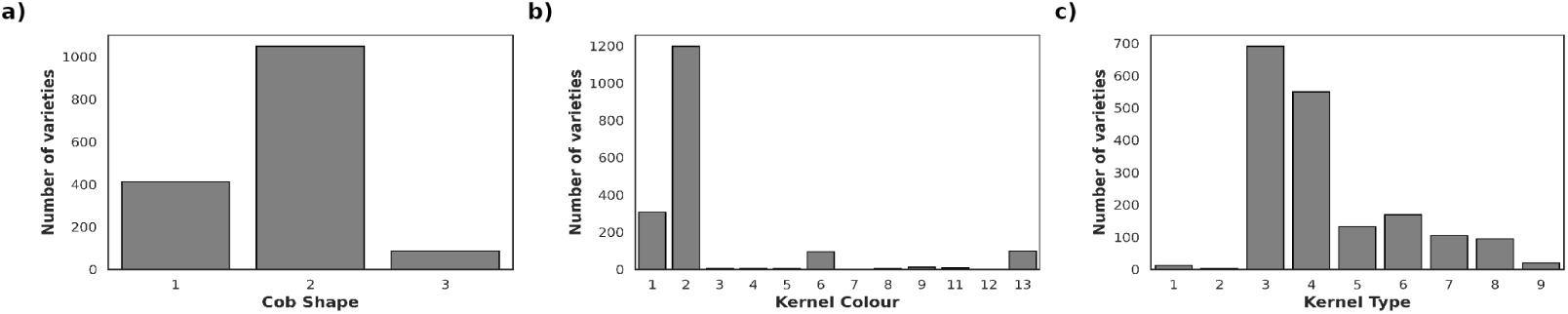
Distribution of notes for three Distinctness, Uniformity, and Stability (DUS) characters from the USDA-NPGS dataset: (a) Cob Shape; (b) Kernel Colour; and (c) Kernel Type.

**Figure 9.**
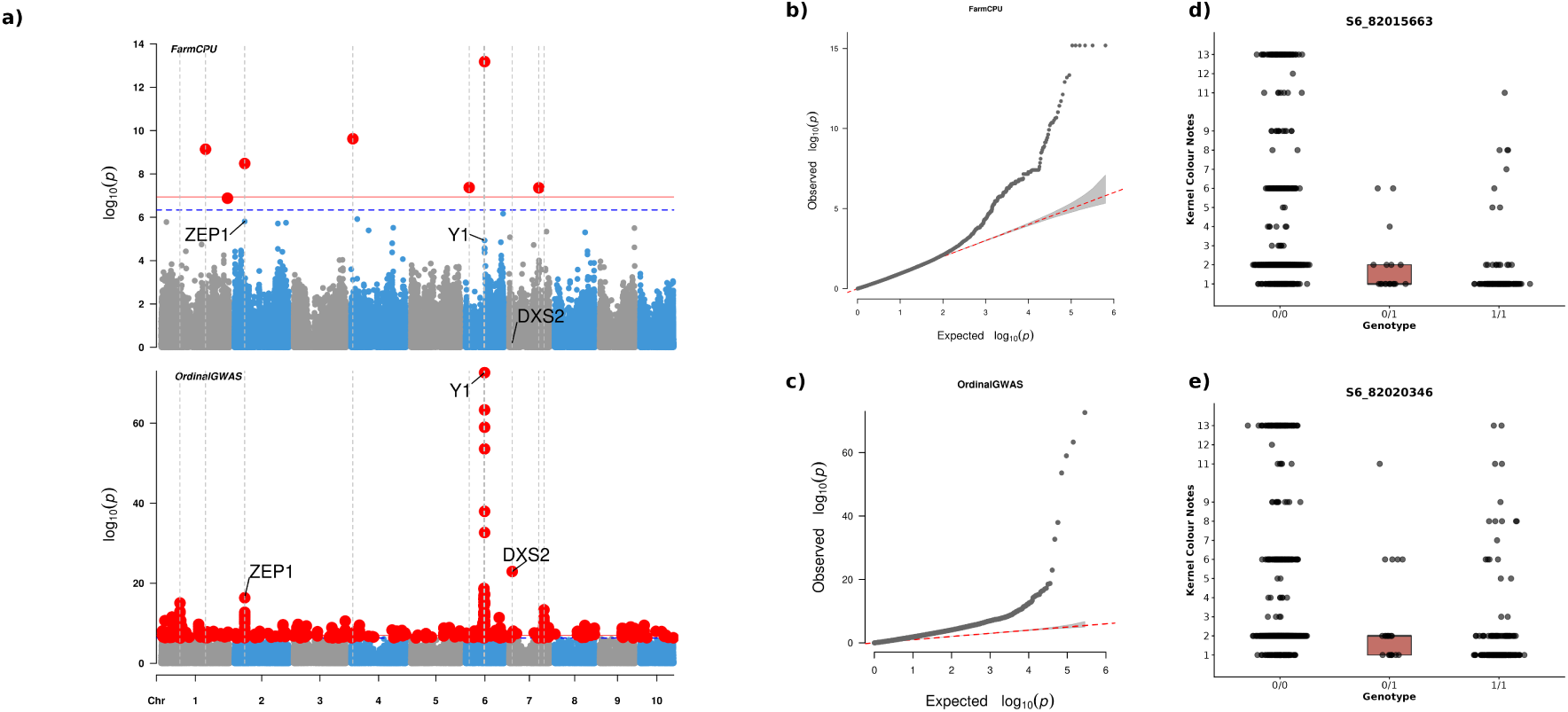
Genome-wide association study (GWAS) of trait “Kernel Colour” using the USDA-NPGS dataset (*n* = 1746). (a) Manhattan plots showing genomic SNP positions versus *p*-values expressed as – *log*_10_(*p*) from the FarmCPU (top) and OrdinalGWAS (bottom) analysis. The red solid line indicate the stringent Bonferroni significance threshold (FWER = 5%, *p* = 5.31 × 10^−8^), while the blue dashed line marks the suggestive threshold (FWER = 20%, *p* = 2.12 × 10^−7^). (b,c) QQ plot comparing observed and expected *p*-value distributions from the FarmCPU (top) and OrdinalGWAS (bottom) analysis. (d,e) Boxplot showing distribution of the “Kernel Colour Notes” across the three genotypic classes at the most significant SNPs from the FarmCPU (top) and OrdinalGWAS (bottom) analysis.

**Figure 10.**
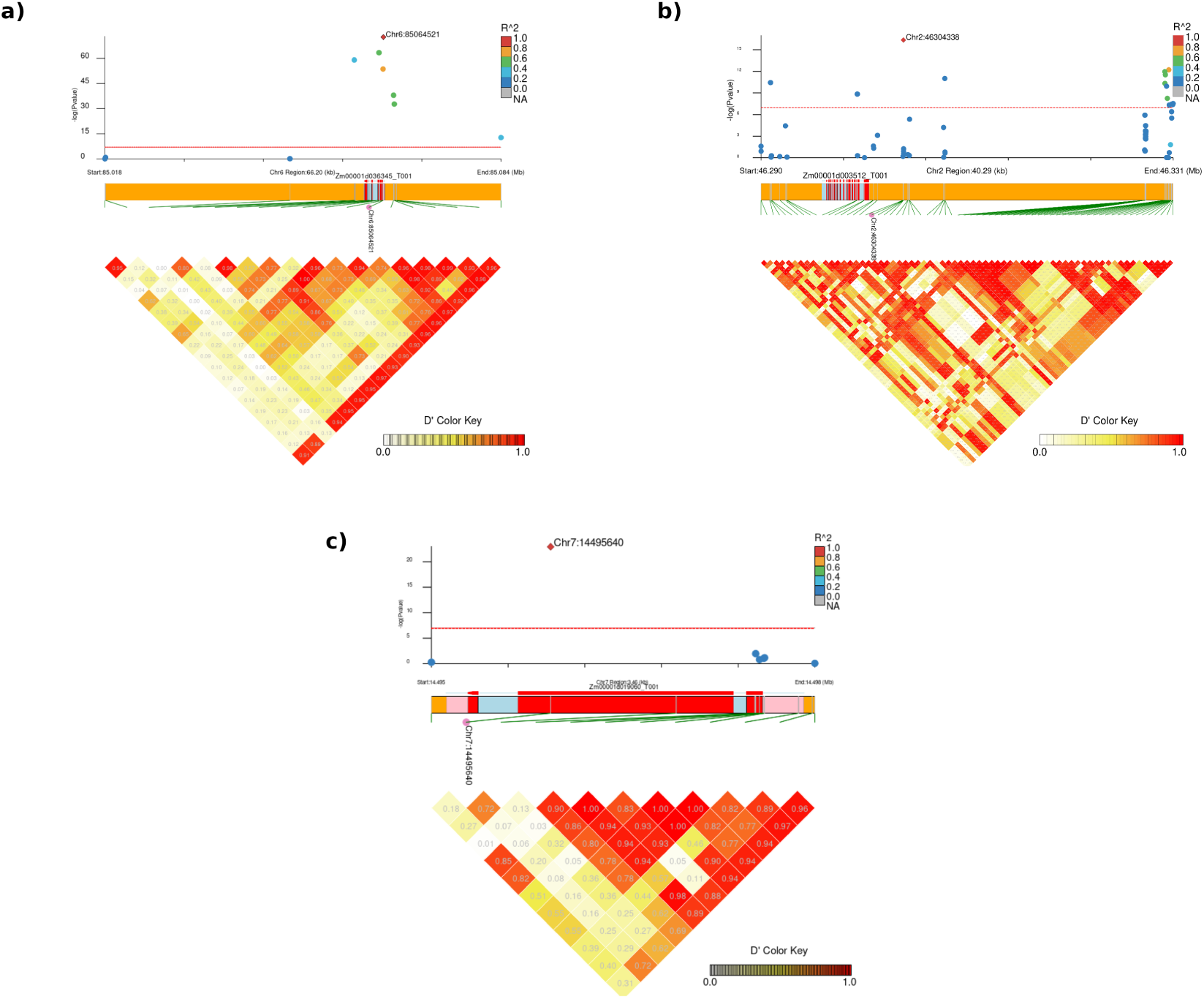
Linkage disequilibrium (LD) structure of three genomic interval on chromosome 2, 6 and 7 with the identified QTL for DUS character “Kernel Colour (KC)”. The Manhattan plot shows a genomic SNP positions versus *p*-values expressed as – *log*_10_(*p*) from OrdinalGWAS, the red dotted line indicate the Bonferroni significance threshold (FWER = 5%; *p* = 5.3 × 10^−8^). The genomic region surrounding the lead-SNP is visualized with its local LD and haplotype block structure. The lead SNP is depicted by a diamond-shaped marker in the upper association track. All neighboring SNPs are color-coded based on their pairwise linkage disequilibrium *r*^2^ with the lead-SNP, indicating the strength of their statistical correlation. The lower panel displays an LD heatmap calculated using *D*^′^, illustrating the historical recombination patterns and local LD architecture.

Although these loci represent biologically plausible candidates underlying variation in kernel colour, alleles at the associated SNPs did not consistently discriminate among all states of expression (i.e. notes), limiting their direct use as diagnostic markers under UPOV BMT Model 1. Local score analysis based on OrdinalGWAS *p*-values (Figure 11) nevertheless identified a QTL interval spanning 3.2 Mb on chromosome 6 that encompassed *Y1*, supporting its role in the genetic control of kernel colour (Supplementary Table 6). In contrast, no QTL intervals were detected for the loci on chromosomes 2 and 7 using this approach.

**Figure 11.**
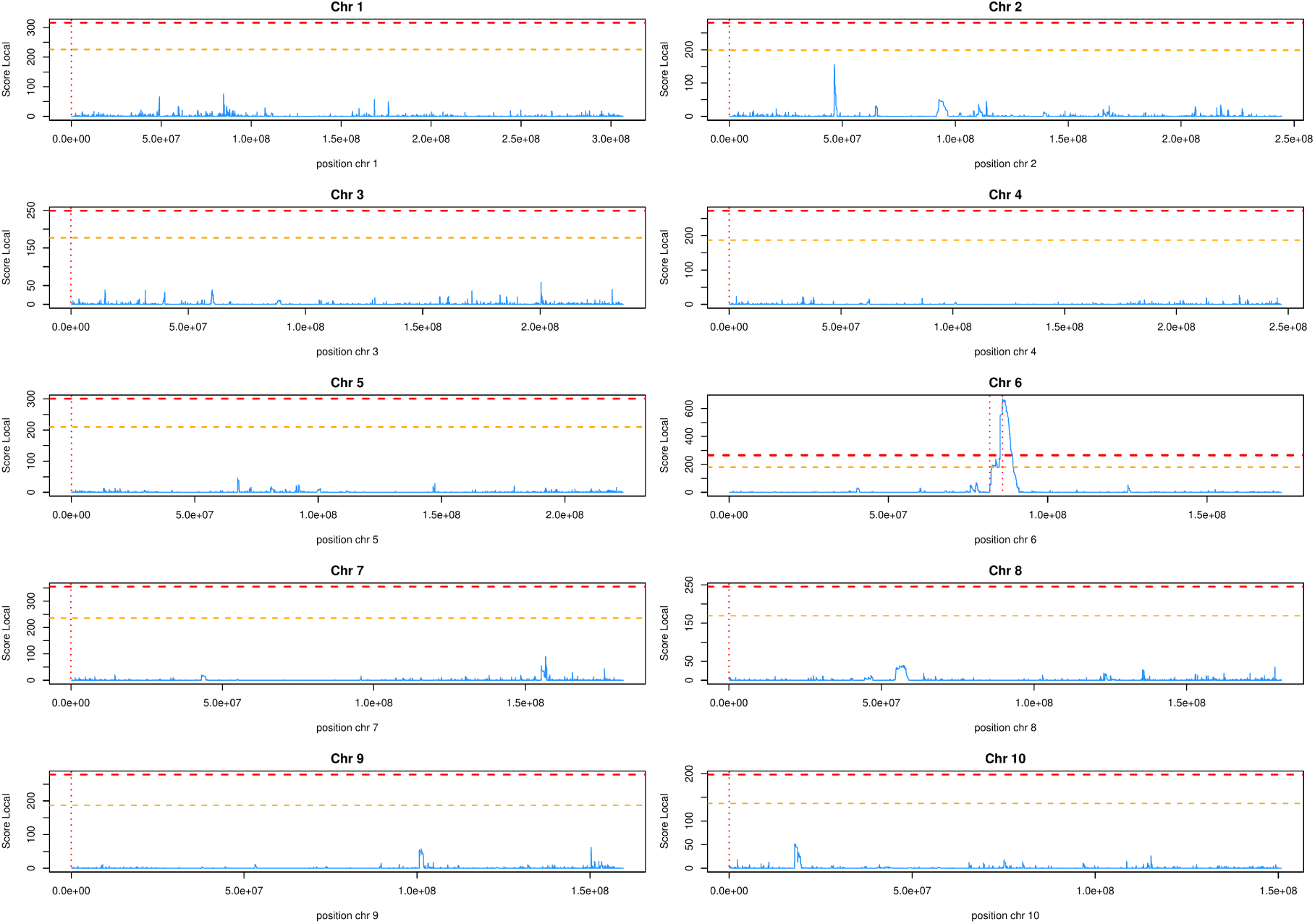
Detection of quantitative trait loci (QTL) associated with the DUS characteristic “Kernel Colour (KC)” using the local score approach. The line plot shows values of the Lindley process (local score) computed from OrdinalGWAS *p*-values using a shape parameter of *ξ* = 1. Horizontal dashed lines indicate genome-wide significance thresholds derived from the Gumbel extreme value distribution at *⍺* = 0.05 (orange) and *⍺* = 0.01 (red). Vertical red lines delineate the boundaries of the detected QTL region.

For the trait ‘kernel type’, which aligns with DUS characteristic “Ear: type of grain” (CPVO 34), genome-wide association analysis identified several significant association peaks; however, none co-localized with genes of known function related to kernel morphology (Supplementary Figure 6 Supplementary Table 5). Similarly, a GWAS for the trait ‘cob shape’, which aligns with “Ear: shape” (CPVO 28) detected multiple associated loci, but no previously characterized candidate genes could be assigned based on current functional annotation (Supplementary Figure 6 Supplementary Table 5).

### Genomic prediction of DUS character states of expression using XGBoost

The predominantly polygenic architecture of most DUS characteristics did not allow the development of single diagnostic molecular markers under UPOV BMT Model 1. We therefore implemented a two-step genomic prediction framework that combines GWAS-based marker selection with XGBoost machine-learning classification (Pudjihartono et al. 2022; Al-Mamun et al. 2025) to predict the notes of DUS characteristics using multiple markers. Within this framework, each DUS characteristic was recoded into binary contrasts between pairs of states of expression (notes), which enables case–control GWAS to identify informative markers for prediction (see Materials and Methods). Logistic regression was used to detect SNPs with significant allele-frequency differences between note categories, and the selected SNPs were subsequently used as predictors in XGBoost models. Prediction models were trained independently for each of the 21 DUS characteristics and evaluated using cross-validation and an independent validation dataset.

Predictive performance and model stability varied across DUS characteristics, as reflected by macro F1 scores obtained across five cross-validation folds. Prediction models for all but three characteristics (CPVO 20, CPVO 22.2, and CPVO 29) showed high stability (cross-validation standard deviation < 0.05), indicating consistent performance across folds (Figure 12 a). Among the 19 characteristics for which models showed stable performance, five achieved high predictive accuracy, with mean macro F1 scores exceeding 0.90: CPVO 34 (0.99), CPVO 7 (0.95), CPVO 4 (0.92), CPVO 14 (0.91), and CPVO 8 (0.90). These characteristics comprised three or fewer states of expression (notes), with the majority of varieties assigned to a single dominant note (Figure 3 Supplementary Table 7). The resulting larger training sample sizes for the predominant class likely contributed to the high classification performance.

**Figure 12.**
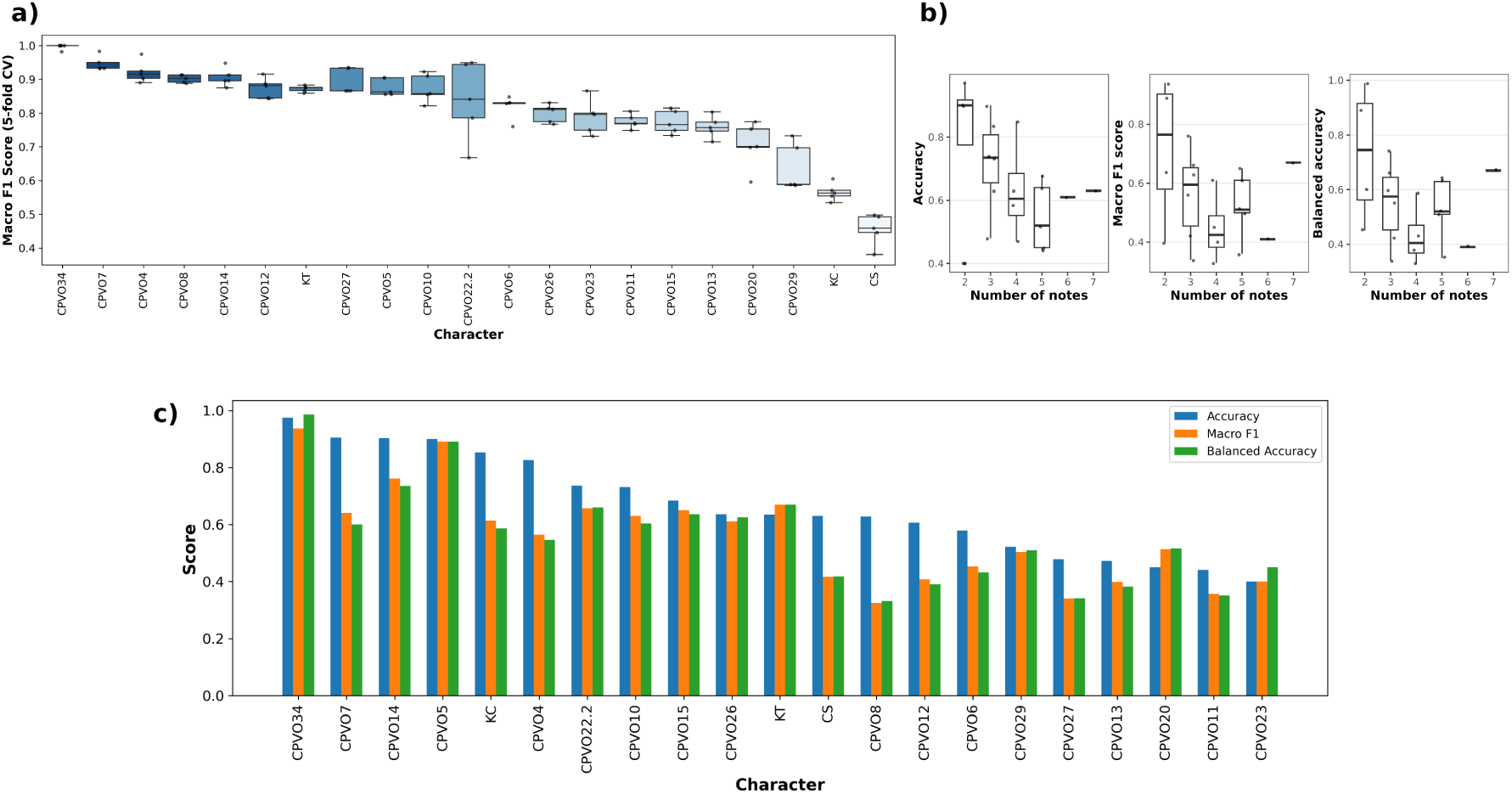
Performance evaluation of XGBoost models for maize DUS character prediction. a) Distribution of macro F1 scores for XGBoost models across five fold cross-validation of distinctness, uniformity and stability (DUS) characters. b) Effect of number of classes (notes) on diverse prediction accuracy matrices. c) Prediction performance of XGBoost models on independent test data for 21 distinctness, uniformity, and stability (DUS) characters, evaluated using three key metrics: Accuracy, Macro F1 score, and Balanced Accuracy. Character IDs correspond to 18 DUS characters from the European dataset as per CPVO testing protocol identifiers, along with three additional characters from the USDA-NPGS dataset: Cob shape, kernel colour, and kernel type.

In contrast, stable models with lower mean macro F1 scores (<0.60), including kernel colour (USDA data) and CPVO 13, were typically associated with characteristics comprising more than three states of expression and more evenly distributed categories, indicating that increased phenotypic complexity reduces classification accuracy. An exception was “Ear: shape” (CPVO 28), which showed low predictive performance despite having only three states of expression with adequate sample sizes (Figure 12a; Supplementary Table 7). Consistent with this pattern, Spearman’s rank correlation revealed a negative relationship between the number of states of expression and predictive accuracy (*ρ* = −0.456, *p* = 0.038; Figure 12b), which was further supported by the Jonckheere–Terpstra test (*p* = 0.030). However, this trend was not observed for the macro F1 (*p* = 0.286) or balanced accuracy (*p* = 0.263) metrics. Neither model stability nor mean macro F1 score correlated with class imbalance, indicating that stratified sampling and the application of SMOTEN effectively mitigated unequal category representation during model training (Figure 3 Figure 12a). Instead, variation in model performance is more plausibly explained by limited sample sizes within validation folds for rare states of expression, as reflected in the distribution of notes across DUS characteristics (Figure 3, Figure 8).

Prediction accuracy metrics (Accuracy, Balanced Accuracy, and Macro F1) obtained from the independent test set varied substantially across the 21 DUS characters and were generally lower than those observed during cross-validation (Figure 12a,c). Across all characters, the mean test accuracy was 0.67, whereas more balanced metrics were markedly lower (Macro F1: 0.56, Balanced Accuracy: 0.56; Figure 12c, Supplementary Table 7). Overall, characteristics with fewer notes, particularly binary traits such as CPVO5 and CPVO34, showed the strongest predictive performance, with Accuracy, Balanced Accuracy, and Macro F1 approaching or exceeding 0.90. Most characters exhibited clear discrepancies between raw Accuracy and more balanced metrics. For example, kernel colour achieved a high overall accuracy (0.85), but its Macro F1 (0.61) and Balanced Accuracy (0.59) were substantially lower (Figure 12c, Supplementary Table 7). These disparities likely originate from imbalanced datasets, where majority classes are predicted more accurately, while minority classes, which comprise only a few samples, show much poorer predictive performance due to insufficient training examples. For kernel colour, among the four classes (1, 2, 6, and 13) included in the training dataset, only classes 1 and 2 had sufficiently large sample sizes; the remaining two classes had very limited representation and exhibited poor predictive performance (Figure 13, Supplementary Figure 7). This pattern indicates that, although the model correctly classified many samples overall, its performance on less frequent classes remained considerably weaker. Similar trends were observed for CPVO8 and CPVO12, where moderate prediction accuracies contrasted sharply with low Macro F1 and Balanced Accuracy values, suggesting a systematic bias toward dominant classes. A related pattern appeared for cob shape: prediction accuracy was correlated with class sample size, yet two of the three classes showed unexpectedly low accuracy despite reasonable sample sizes (Figure 12c, Figure 13).

**Figure 13.**
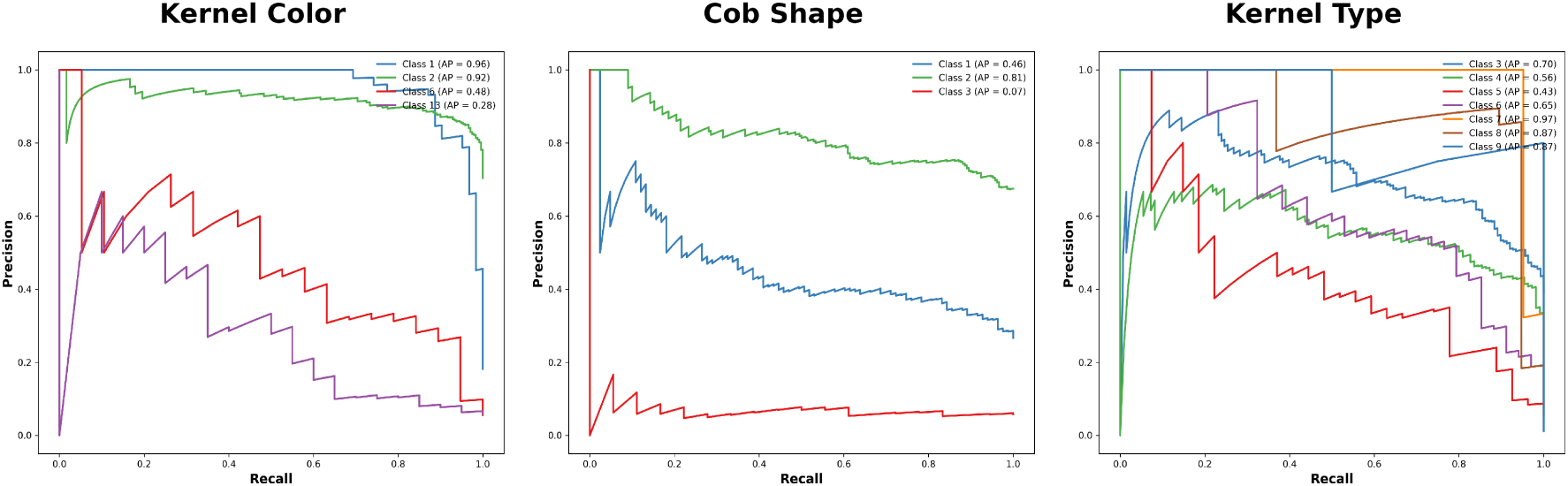
Precision-recall curves showing classification performance of model for each class (note) of “Cob Shape”, “Kernel Type” and “Kernel Colour” characters (USDA-NPS). The Average Precision (AP) summarizes overall performance for each class.

In contrast, for kernel type, prediction accuracy did not correlate with class sample size. Surprisingly, the two most prevalent classes (3 and 4) showed relatively poor prediction accuracy, whereas the less frequent classes 7, 8, and 9 were predicted with substantially higher precision (Supplementary Figure 7, Figure 13). Kernel type with seven notes achieved moderate accuracy alongside high Balanced Accuracy and Macro F1 values, demonstrating effective performance despite the large number of notes.

## Discussion

We analyzed historical DUS phenotypic data from 348 registered maize hybrid varieties examined by four major European examination offices (EOs) and generated a comprehensive genome-wide marker dataset comprising more than 460,000 SNPs using high-density SNP arrays (600K) and low-coverage whole-genome sequencing (lcWGS). This integrated resource enabled an evaluation of molecular markers in maize DUS testing within the framework of UPOV BMT Models 1 (“Charac-teristic-Specific Molecular Markers”) and 2 (“Combining phenotypic and molecular distances in the management of variety collections”). Marker density is an order of magnitude higher than previous studies in maize or other crops (Cockram et al. 2012; Thomasset et al. 2015; Achard et al. 2020; Yang et al. 2021; Liu et al. 2022; Abd El-Wahab et al. 2023; Zanella et al. 2026). A high marker density is advantageous as marker number determines the successful application of UPOV BMT models (Cockram et al. 2012).

### Implications of genetic structure for DUS testing

Principal component analysis identified distinct population clusters among European maize varieties, broadly corresponding to their examination offices (AGES, BSA, GEVES, NEBIH). These clusters indicate that differences in breeding practices, selection criteria, and regional adaptation have contributed to shaping the genetic structure of European maize. A reduced diversity within an EO portfolio may increase the difficulty of distinguishing closely related varieties based solely on morphological DUS characteristics, potentially complicating phenotypic distinctness assessments. The observed population structure has important implications for future studies aimed at development of new DUS testing framework as well as in the practical application of UPOV BMT models. Under Model 1, which requires character-specific diagnostic markers, strong genetic stratification may lead to population-specific marker–trait associations. For example, anthocyanin-related traits in barley are controlled by loci such as *ant1* and *ant2* that segregate in winter but not in spring varieties, illustrating that causal variants for a given DUS trait may be present only in particular genetic subgroups (Yang et al. 2021). If reference collections do not adequately represent the underlying genetic diversity, marker effects may not be transferable across groups, potentially affecting the reliability of marker-based distinctness assessments (Reyna and Sneller 2001). Therefore, extensive validation of candidate markers across a broad and representative range of varieties will be essential to ensure robustness and general applicability (Arens et al. 2010; Platten et al. 2019). These findings emphasize that the development of molecular markers linked to specific states of expression (notes) must be accompanied by careful selection of reference varieties representing the full spectrum of phenotypic variation and genetic backgrounds across examination offices. Such representation is critical to ensure that candidate varieties are assessed against an appropriately diverse reference collection and that marker-based approaches support unbiased and consistent DUS examination.

Implementation of UPOV BMT Model 2, which supports the management of reference collections through molecular similarity measures, may likewise be affected by underlying genetic structure ((CPVO) 2011). Genetic distances within and between population clusters do not necessarily correspond to phenotypic distinctness based on DUS characteristics ((CPVO) 2011). Consequently, predictive thresholds or similarity criteria may require validation across different genetic groups or examination offices to ensure consistent and reliable application and to avoid erroneous assessments of phenotypic similarity.

### Genetic architecture of DUS characters limits UPOV BMT Model 1 implementation

GWAS analysis revealed that most DUS characters are polygenic, with at least five significant loci detected for each character except “tassel: anthocyanin coloration of glumes excluding base (CPVO8)”. This polygenic architecture was confirmed using a larger public USDA-NPGS maize panel and also aligns with findings in rice, barley, and fenugreek (Yang et al. 2021; Liu et al. 2022; Abd El-Wahab et al. 2023). Critically, none of the GWAS-detected SNPs showed sufficient discriminatory power to differentiate between notes of any character, consistent with previous studies in other crops (Jones and Mackay 2015; Yang et al. 2021; Liu et al. 2022). This lack of discriminatory power creates significant challenges for UPOV BMT Model 1 implementation, which requires verified associations between molecular markers and specific DUS characters (UPOV 2011; Yang et al. 2021). This limitation likely reflects incomplete linkage disequilibrium between associated SNPs and the underlying causal variants, as well as the restricted ability of bi-allelic SNP markers to capture the full spectrum of functional genetic variation. A comparable situation was reported in a recent study in pea (*Pisum sativum*) investigating the genetic architecture of seven classical Mendelian characters using GWAS (Feng et al. 2025). Although broad association peaks were identified, these signals did not directly resolve the causal polymorphisms. Subsequent fine-mapping approaches, including bulk-segregant analysis, KASP genotyping, and recombination breakpoint mapping, revealed that the causal variants comprised large upstream deletions, transposon insertions, and small coding insertions or deletions, which were only imperfectly tagged by nearby bi-allelic SNP markers.

Therefore, for an application of our marker-characteristic associations, fine-mapping of GWAS loci is required to identify causal polymorphisms and to develop allele-specific markers suitable for reliable prediction of states of expression (notes) of DUS characteristics. This approach is particularly relevant for characteristics controlled by major-effect genes. Successful examples have been reported in barley for *VERNALIZATION-H1* (*VRN-H1*) and *VRN-H2* (Cockram et al. 2007, 2009; Yang et al. 2021; Cosenza et al. 2024), and in tomato for disease resistance genes (Arens et al. 2010; UPOV 2020), where functional polymorphisms enabled the development of diagnostic markers. Advances in high-resolution genomics, including pan-genome resources, now facilitate more rapid fine-mapping of multiple QTL and improve the detection of structural and presence–absence variation underlying phenotypic differences (Zanini et al. 2022; Li and Zhou 2025). While these examples focus on Mendelian or major-effect genes, the polygenic nature of most DUS traits suggests that a strict one-to-one marker-trait relationship is often unattainable. In such cases, the utility of UPOV BMT Model 1 might be extended by identifying stable combinations of markers that collectively reach the required threshold of discriminatory power, as demonstrated previously (Cockram et al. 2007) or in the present study.

Our identification of biologically plausible candidate genes illustrates how functional annotation can guide marker development for potential application under UPOV BMT Model 1. For the DUS characteristic “Tassel: anthocyanin coloration of glumes excluding base (CPVO 8)”, we identified a likely candidate gene: *C1* is a MYB transcription factor that regulates key genes of the anthocyanin biosynthesis pathway (*A1*, *A2*, *Bz1*, *Bz2*, and *Pr1*) controlling tissue-specific pigmentation (Cone et al. 1986; Sainz et al. 1997). The *C1* locus is characterized by extensive allelic diversity, largely driven by structural variation in regulatory regions, including insertions and excisions of transposable elements such as *Spm* (*Suppressor–mutator*) and *Ds* (*Dissociation*) (Paz-Ares et al. 1990; Singer et al. 1998). Reported alleles include the functional wild-type *C1* conferring purple pigmentation, the loss-of-function *c1* allele, the dominant inhibitor *C1-I*, the partially compensatory *C1-S*, and mutable *c1-m* alleles associated with somatic instability. Such well-characterized allelic variation provides a robust biological foundation for the development of allele-specific markers capable of discriminating relevant states of expression for CPVO 8 in DUS examination (Paz-Ares et al. 1990; Singer et al. 1998).

Beyond established loci, several novel candidate genes were also identified. Notably, for “Tassel: curvature of lateral branches (CPVO11)”, the Agamous-like MADS-box protein AGL16 emerged as a new candidate gene. Given that maize ZAG/ZMM family MADS-box genes are critical for floral organ identity and meristem determinacy, AGL16 may integrate into higher-order regulatory complexes. Such interactions likely modulate inflorescence meristem architecture, indirectly influencing branch initiation, length, and displacement angle (Danilevskaya et al. 2008; Zhao et al. 2021). Overall, our findings demonstrate that for a subset of DUS characteristics controlled by major-effect genes, candidate gene identification provides concrete targets for the development of diagnostic markers. Such markers represent a prerequisite for reliable implementation of UPOV BMT Model 1, provided that their specificity and stability are validated across diverse genetic backgrounds.

### Genomic prediction supports UPOV BMT Model 2 implementation

The two-step genomic prediction (GP) framework, combining GWAS-based marker selection with XGBoost classification, was designed to predict states of expression (notes) of DUS characteristics in support of UPOV BMT Model 2, which allows the use of molecular information for reference collection management without replacing phenotypic DUS examination. Across 21 characteristics, the approach achieved a mean prediction accuracy of 0.67, exceeding values reported for rice using rrBLUP (0.48) (Liu et al. 2022). This improved performance likely reflects methodological advantages. Linear models such as GBLUP and rrBLUP assume continuous phenotypes and additive marker effects, which are not fully compatible with ordinal DUS scoring (Kizilkaya et al. 2014; Merrick et al. 2022). In contrast, XGBoost captures non-linear relationships and marker interactions and accommodates imbalanced distributions of states of expression through stratified sampling and oversampling strategies (Montesinos-López et al. 2015; Montesinos-López et al. 2022). Most characteristics showed high cross-validation stability, supporting the robustness of the approach. XGBoost has also been shown to outperform linear GWAS models for categorical trait prediction in quinoa (*Chenopodium quinoa*) (Sandell et al. 2024).

Prediction accuracy on independent test data varied among characteristics, in line with previous studies in rice and barley (Jones and Mackay 2015; Liu et al. 2022; NIAB 2025). This variation likely reflects differences in genetic architecture, heritability, and the number and distribution of states of expression. Within a Model 2 context, such models are intended to support the identification of varieties with similar phenotypic profiles and to guide the selection of appropriate comparators within structured reference collections. In combination with the observed population structure among examination offices, the results indicate that genomic prediction can provide complementary decision-support information, while phenotypic assessment remains the basis for distinctness determination. Prediction performance was primarily determined by (i) the number of states of expression and (ii) the number of observations per note. Accuracy declined with increasing numbers of notes, consistent with general observations that multi-class prediction becomes more challenging as class number and granularity increase (Sun et al. 2021). This likely reflects increased overlap among adjacent notes and unequal representation of states of expression. Rare notes were consistently predicted with lower accuracy, resulting in reduced balanced accuracy and macro F1 values, whereas well-represented notes achieved higher predictive performance. These results indicate that reliable application in a Model 2 framework depends on sufficiently large and balanced training populations. In practical DUS examination, where reference collections may not fully span the range of notes, predictive performance can therefore vary across characteristics and states of expression. Deviations from this general pattern highlight the influence of trait properties. For example, “Ear: shape” (CPVO 28) showed only moderate predictive performance despite few notes and balanced representation, consistent with characteristics that may exhibit limited discriminating power due to measurement variability, observer subjectivity, genotype-by-environment interaction, or restricted phenotypic contrast (Law et al. 2011). In contrast, “Ear: type of grain” (CPVO 34) showed comparatively high predictive performance despite seven notes, likely reflecting simpler genetic control and limited environmental influence. As in conventional DUS testing, careful selection of characteristics with stable expression and high discriminatory power remains essential for marker-based applications.

Overall, robust genomic prediction of DUS characteristics appears feasible when characteristics show stable expression, relatively simple genetic control, and adequate representation of all states of expression. Under these conditions, high concordance between predicted and observed notes can be achieved. Within a UPOV BMT Model 2 framework, such concordance may support reference collection management by reducing unnecessary phenotypic comparisons, while maintaining phenotypic assessment as the basis for distinctness decisions. Earlier studies in barley reported limited success due to lower prediction accuracy and weaker correspondence between predicted and observed notes (Jones and Mackay 2015). More recent approaches based on multi-year data have demonstrated the potential to integrate genomic prediction into currently used Combined Over Years Distinctness (COYD)-like frameworks (Roberts et al. 2026). A comparable extension of the XGBoost-based framework could be envisaged, provided that multi-year, multi-plant DUS datasets are available and predictive performance is rigorously validated.

### Study limitations and methodological challenges

Both the identification of marker–trait associations for DUS characteristics and the genomic prediction of states of expression (notes) were influenced by characteristics of the available datasets. A relatively high proportion of missing observations for some characteristics limited the construction of common training and independent test sets across traits (Yang et al. 2021; Zanella et al. 2026). As a result, character-specific datasets were used, which restricted unified external validation and direct comparison of marker-based predictions with classical phenotype-based approaches such as COYD testing (Cockram et al. 2015; Yang et al. 2021). Similarly, the absence of common validation sets precluded prediction for entirely independent varieties and the calculation of genomic-derived phenotypic distances for direct comparison with observed DUS outcomes.

Our study utilized highly heterozygous F1 hybrid varieties rather than on homozygous parental lines, which may reduce power to detect marker–trait associations and affect inference of genetic architecture and prediction accuracy (Technow et al. 2012; Bernardo 2016). The composition of the variety panel was also influenced by constraints, including seed availability (particularly for older varieties), breeder consent, and the absence of unsuccessful candidate varieties, potentially influencing the representation of notes and genetic backgrounds. These factors highlight constraints of historical DUS datasets for molecular analyses. Differences in testing protocols among examination offices (EOs) limited overlap of DUS characteristics and required harmonization of data. Uneven note distributions reduced statistical power, and the lack of original continuous measurements prior to categorical conversion constrained analytical resolution and quantitative modeling. Limited replication across trials also restricted estimation of heritability and environmental effects, which are important for selecting robust DUS characteristics. Overall, these factors reflect the historical design of DUS datasets rather than analytical shortcomings. The present results should therefore be regarded as a proof-of-concept demonstrating the feasibility of integrating molecular markers into maize DUS examination, and highlight priorities for future harmonized and multi-environment DUS data collection (Cockram et al. 2015; Yang et al. 2021).

### Implications for Maize DUS Examination

Across the analyzed dataset, 18 robust QTL were identified for 12 DUS characteristics, together with biologically plausible candidate genes. These results provide a concrete basis for further evaluation of molecular markers under UPOV BMT Model 1, particularly for characteristics controlled by major-effect loci. For example, “Tassel: anthocyanin coloration of glumes excluding base” (CPVO 8) and kernel colour (CPVO 36) yielded associations with well-characterized pigmentation genes, consistent with previous examples of marker use in maize and other crops reported in UPOV documentation. For such characteristics, targeted fine-mapping to resolve causal polymorphisms and the development of allele-specific markers may enable reliable discrimination of relevant states of expression (notes). Validation across diverse genetic backgrounds and reference collections will be essential to ensure marker robustness and transferability.

For the majority of DUS characteristics, which exhibit polygenic control, implementation under UPOV BMT Model 2 appears more immediately feasible. The XGBoost-based genomic prediction framework demonstrated stable and moderate-to-high predictive performance across multiple characteristics, supporting its use as a decision-support tool for reference collection management. Under Model 2, the objective is not to replace phenotypic examination but to improve efficiency by identifying varieties with similar predicted DUS profiles and reducing unnecessary pairwise comparisons.

Key next steps for Model 2 implementation include: (i) training prediction models on expanded, preferably multi-year and replicated DUS datasets; (ii) validating correlations between genomically predicted and observed note-based distances to exceed the required threshold (*r* > 0.6), particularly around the one- to two-note distinctness threshold; and (iii) calibrating genomic similarity thresholds that minimize the risk of excluding truly similar varieties. Validation within established maize reference collections specific for each EO will further ensure consistent application across examination offices.

Overall, this study provides an empirical foundation for integrating molecular information into maize DUS examination. The combination of high-density genome-wide markers, candidate gene identification for selected characteristics, and validated genomic prediction models offers complementary pathways for future implementation of UPOV BMT Models 1 and 2, supporting more efficient and data-informed DUS procedures while maintaining the primacy of phenotypic assessment.

## Author Contributions

A.D.: Conceptualization, Methodology, Investigation, Formal analysis, Software, Writing – Original Draft, Visualization, Project administration. C.H.: Formal analysis, Data curation, Writing – Review & Editing. M.P.: Data curation, Provision of Data, Writing – Review & Editing. A.R.: Data curation, Provision of Data, Writing – Review & Editing. P.S.: Data curation, Provision of Data, Writing – Review & Editing. C.S.: Data curation, Provision of Data, Writing – Review & Editing. K.S.: Conceptualization, Supervision, Funding acquisition, Resources, Writing – Review & Editing.

## Data Availability

In accordance with the grant agreement and consortium contract among the project partners, raw genotypic data and DUS states of expression (notes) cannot be made publicly available, to safeguard the commercial interests of participating breeding partners. Aggregated summary statistics are provided where permissible.

The USDA phenotypic and genotypic datasets used in this study are publicly available. In addition, the R scripts and computational pipelines used for the analysis of genetic and phenotypic variation in both the European maize hybrid panel and the USDA dataset have been deposited in the Zenodo repository (DOI: 10.5281/zenodo.20610279).

## Acknowledgements

We thank Vanessa Hasender for technical help in the genotyping and the members of the INVITE consortium as well as the research group of K. S. for discussions and advice. We thank the Federal Plant Variety Office (Bundessortenamt, BSA) of Germany for curation of DUS data, providing DUS characteristics and seed material for genotyping of the 27 maize varieties used in this study.

