## Supplementary Information for "Leveraging genome-wide association studies and genomic prediction for distinctness, uniformity, and stability (DUS) testing in maize"

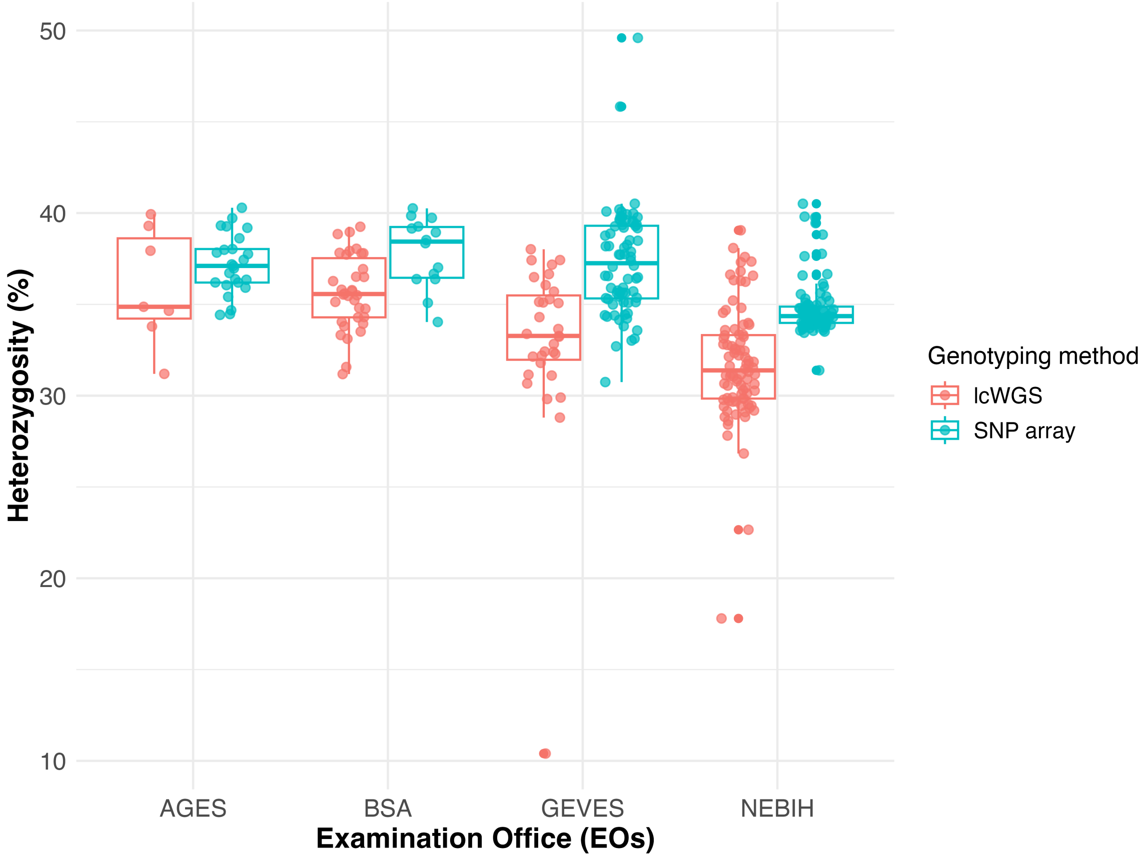


*Supplementary Figure S 1.* Comparison of heterozygosity across different Examination Offices (EOs) observed with two different genotyping methods: low coverage whole genome sequencing (lcWGS) and Axiom 600k SNP genotyping array.

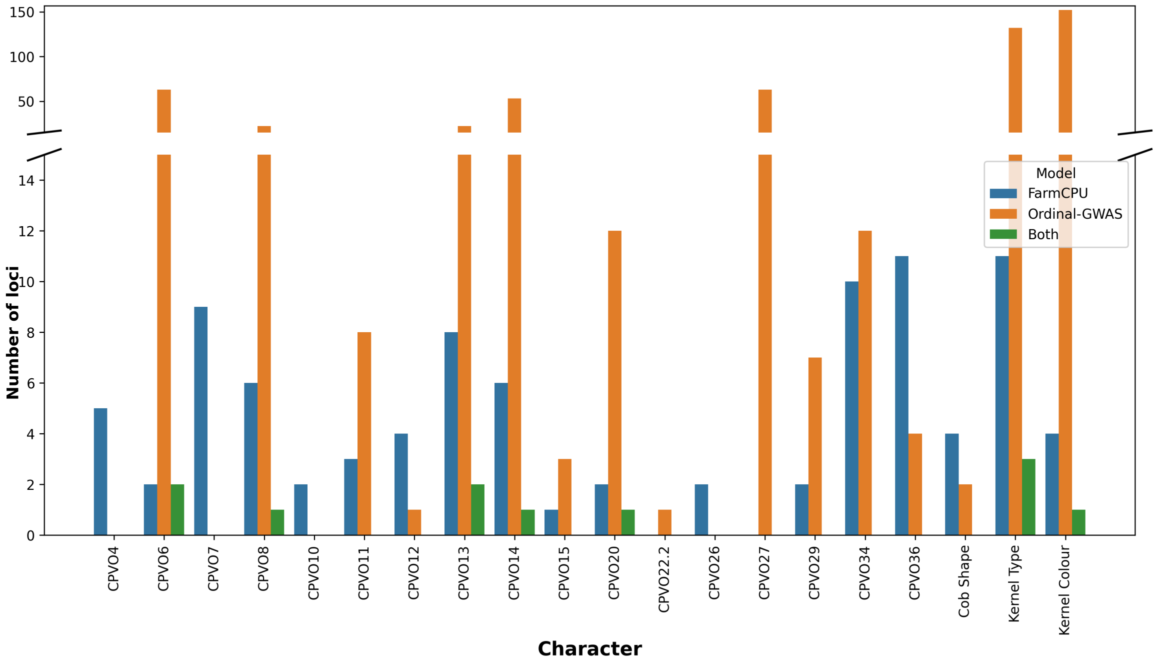


*Supplementary Figure S 2.* Numbers of genomic regions significantly associated DUS trait notes in genome-wide association studies (GWAS) using OrdinalGWAS and FarmCPU methods. The Bonferroni-corrected genome-wide significance thresholds (family-wise error rate, FWER = 5%; $p = 1.16 \times 10^{-7}$ for INVITE and $p = 5.31 \times 10^{-8}$ for USDA-NPGS) was used for determining the significant regions.

### CPVO6

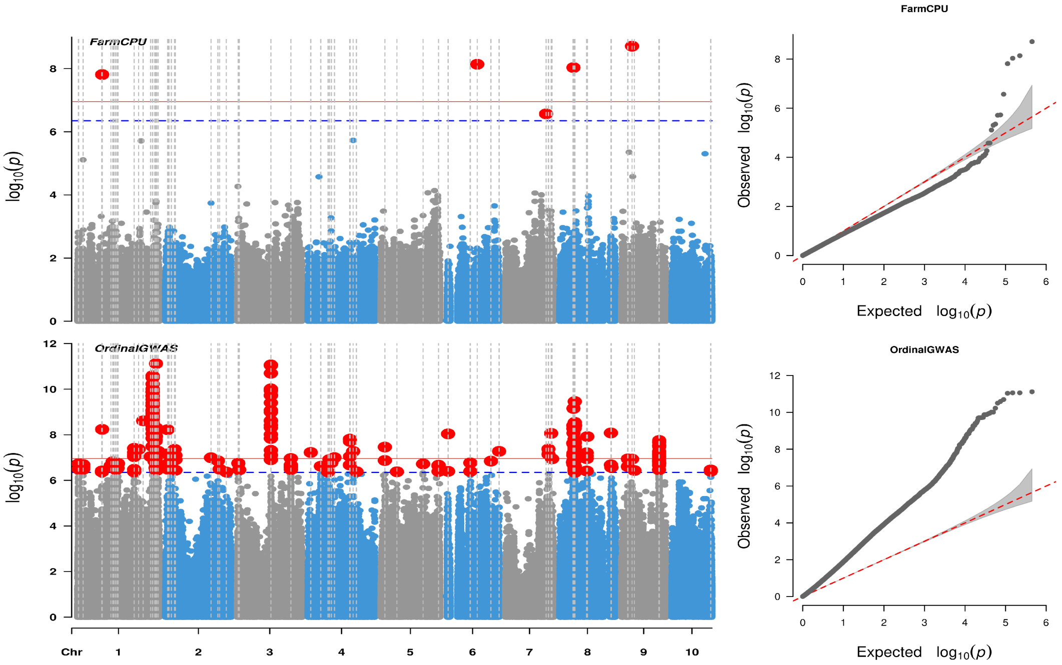


### CPVO10

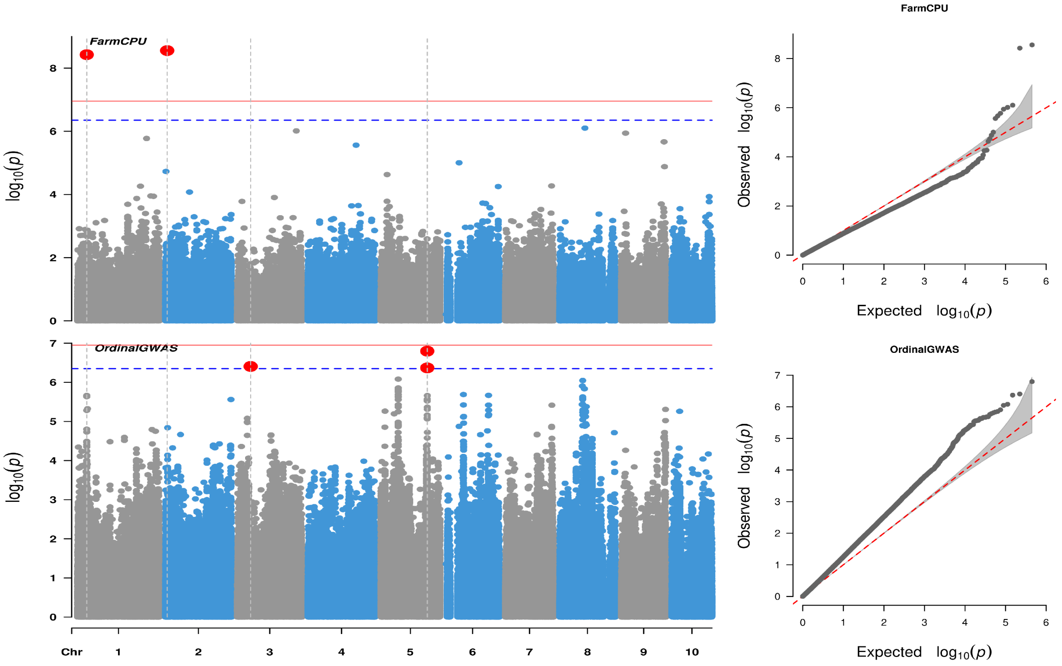

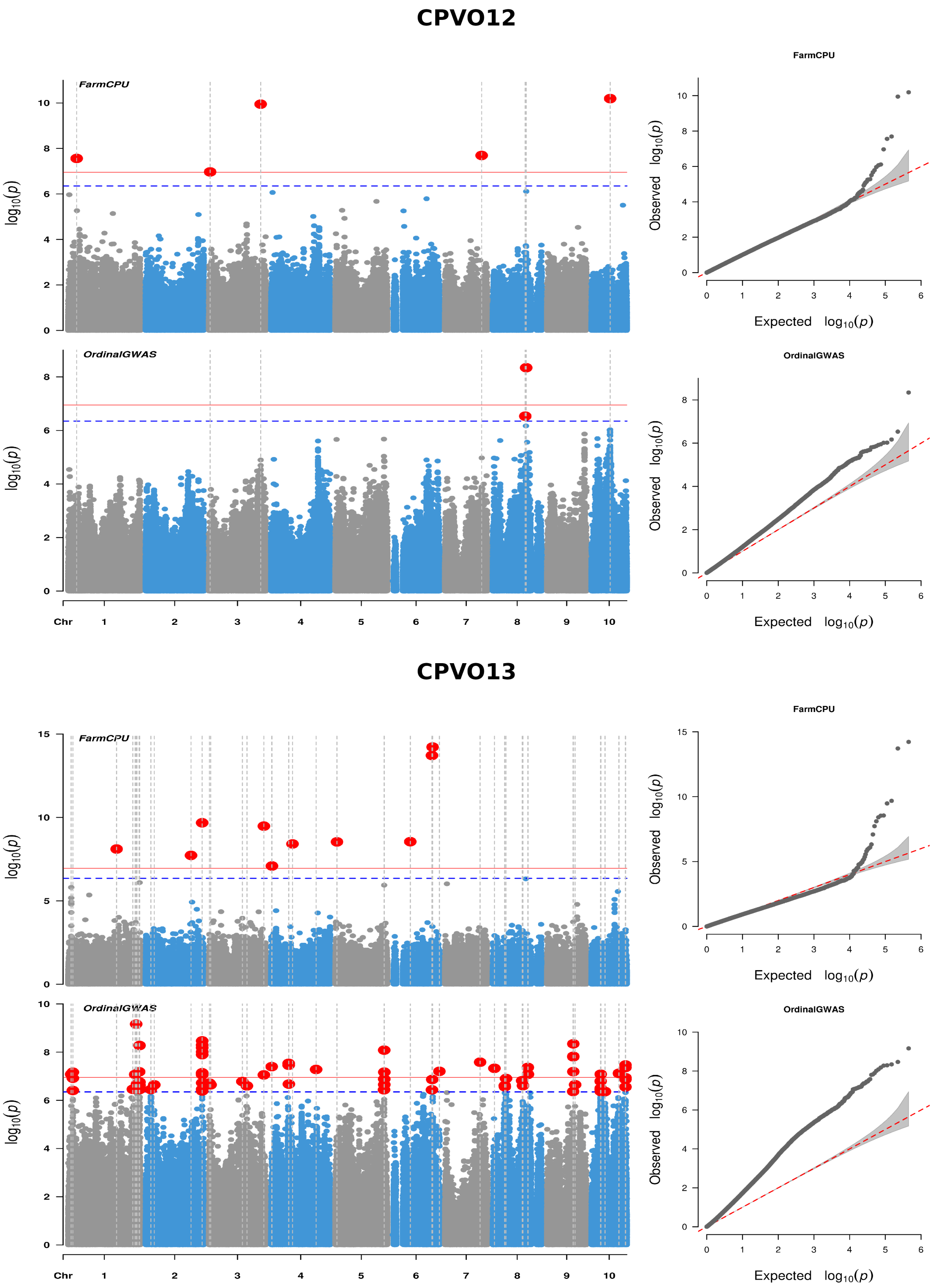


*Supplementary Figure 3.* Genome-wide association study (GWAS) analysis of four high-confidence DUS characters: Tassel: time of anthesis (CPVO6); Tassel: angle between main axis and lateral branches (CPVO10); Tassel: number of primary lateral branches (CPVO12); and Ear: time of silk emergence (CPVO13). In each Manhattan plots show genomic SNP positions versus $p$-values expressed as $-log_{10}(p)$ from the FarmCPU (top) and OrdinalGWAS analysis (bottom). The red solid line indicate the stringent Bonferroni significance threshold (FWER = 5% / $p = 1.12 \times 10^{-7}$), while the blue dashed line marks the suggestive threshold (FWER = 20% /$p = 4.47 \times 10^{-7}). The vertical dashed lines depict SNPs which are significant (FWER = 5% / $p = 1.12 \times 10^{-7}$) with at least one of the two methods. The accompanying QQ plots compare observed and expected $p$-value distributions from the FarmCPU and OrdinalGWAS analysis.

### CPVO4

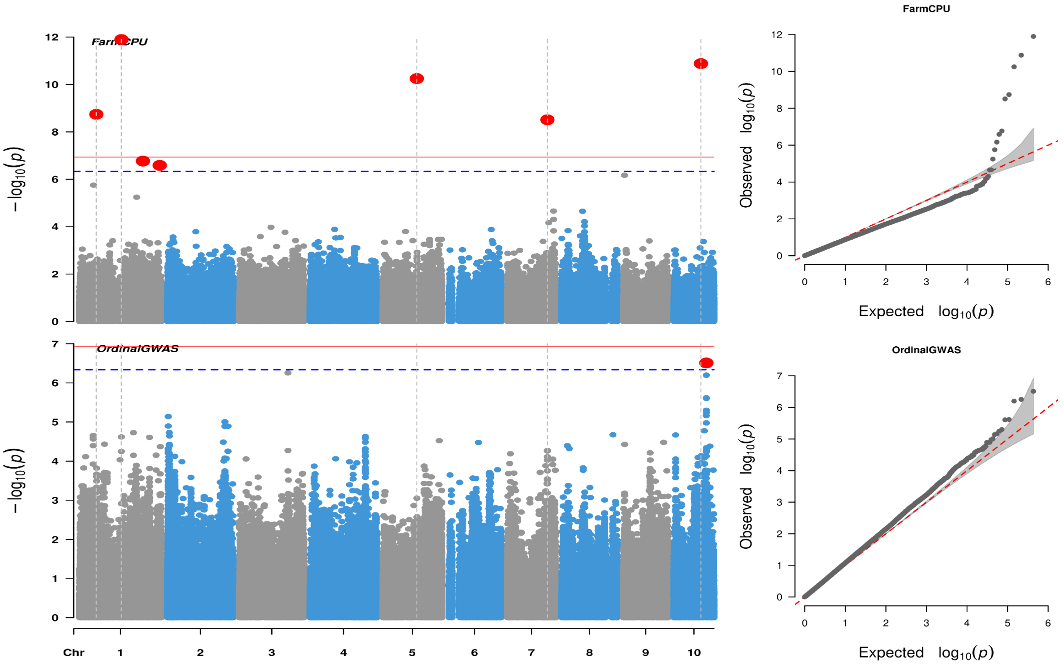
FarmCPU
OrdinalGWAS
$-\log_{10}(p)$
Chr 1 2 3 4 5 6 7 8 9 10
Observed $\log_{10}(p)$
Expected $\log_{10}(p)$

### CPVO7

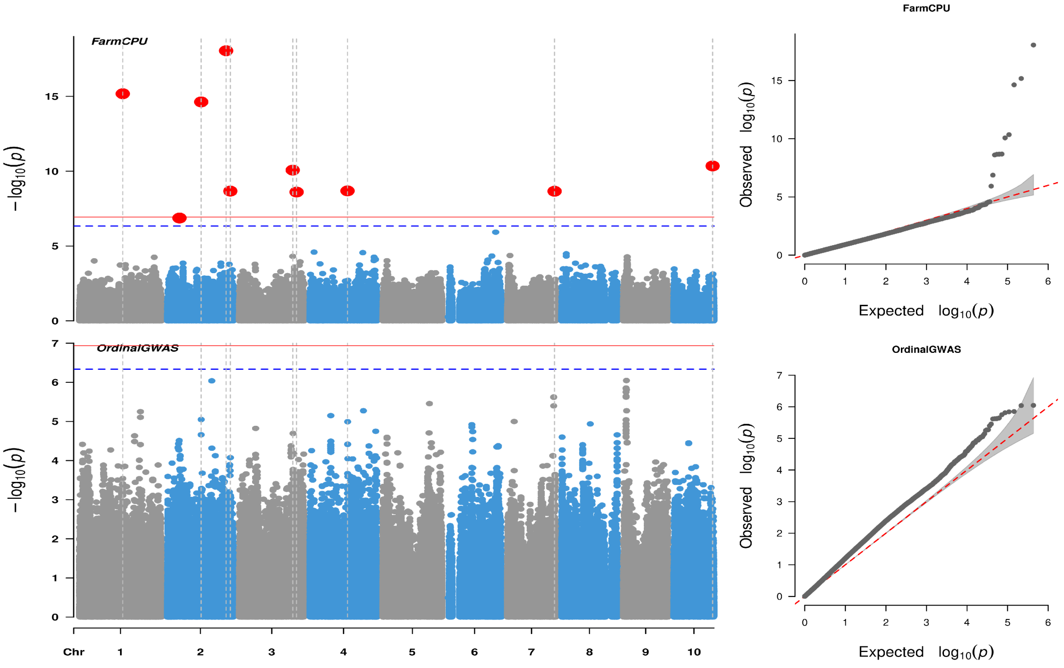
FarmCPU
OrdinalGWAS
$-\log_{10}(p)$
Chr 1 2 3 4 5 6 7 8 9 10
Observed $\log_{10}(p)$
Expected $\log_{10}(p)$

**CPVO11**

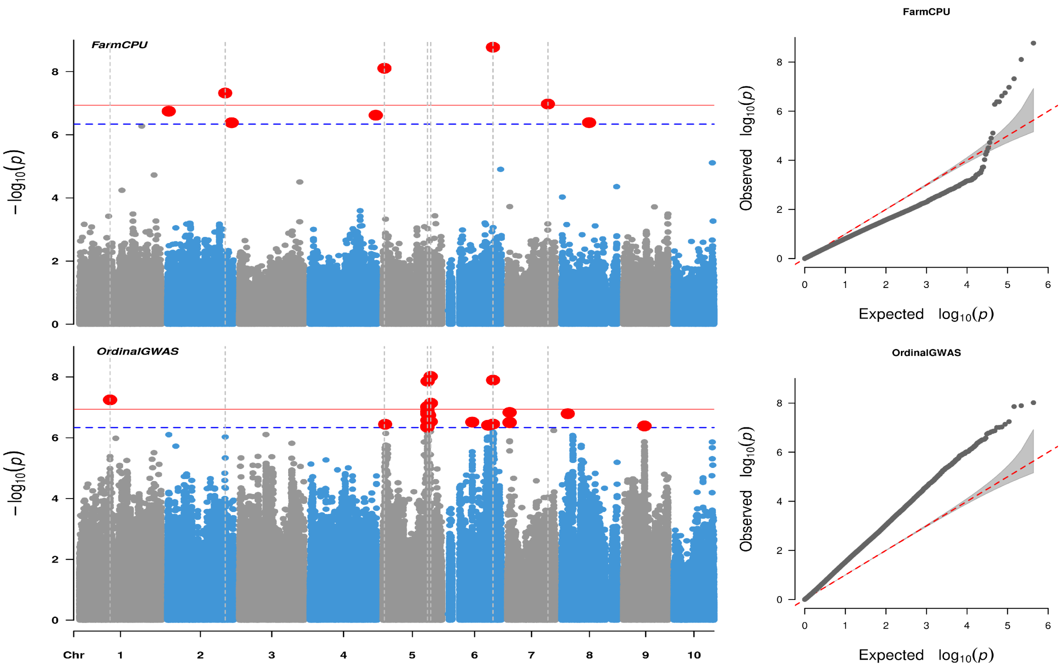


**CPVO14**

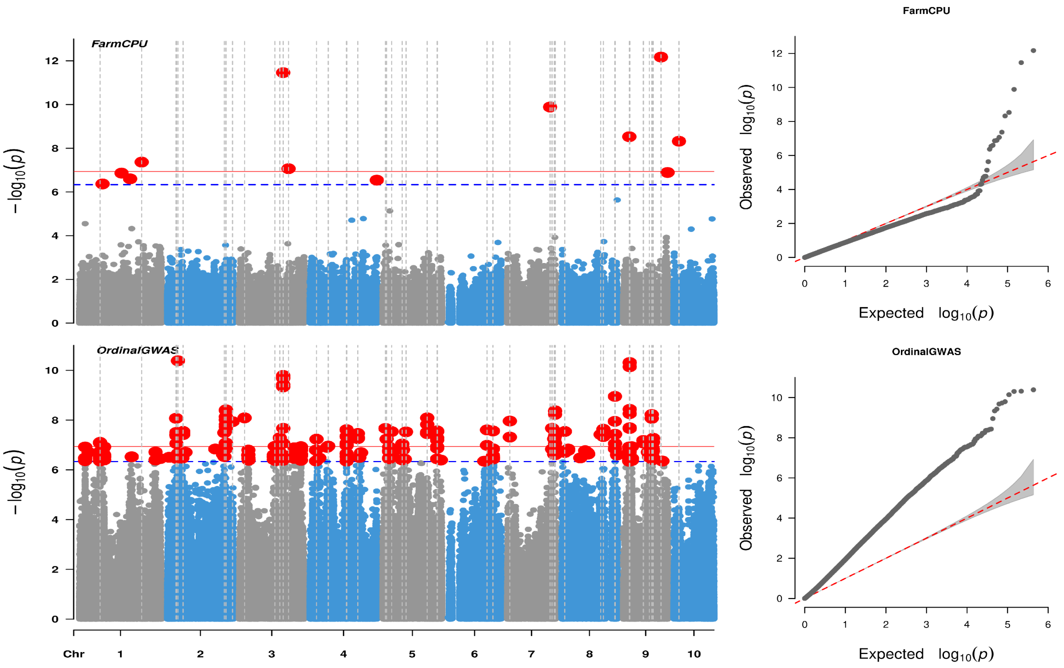

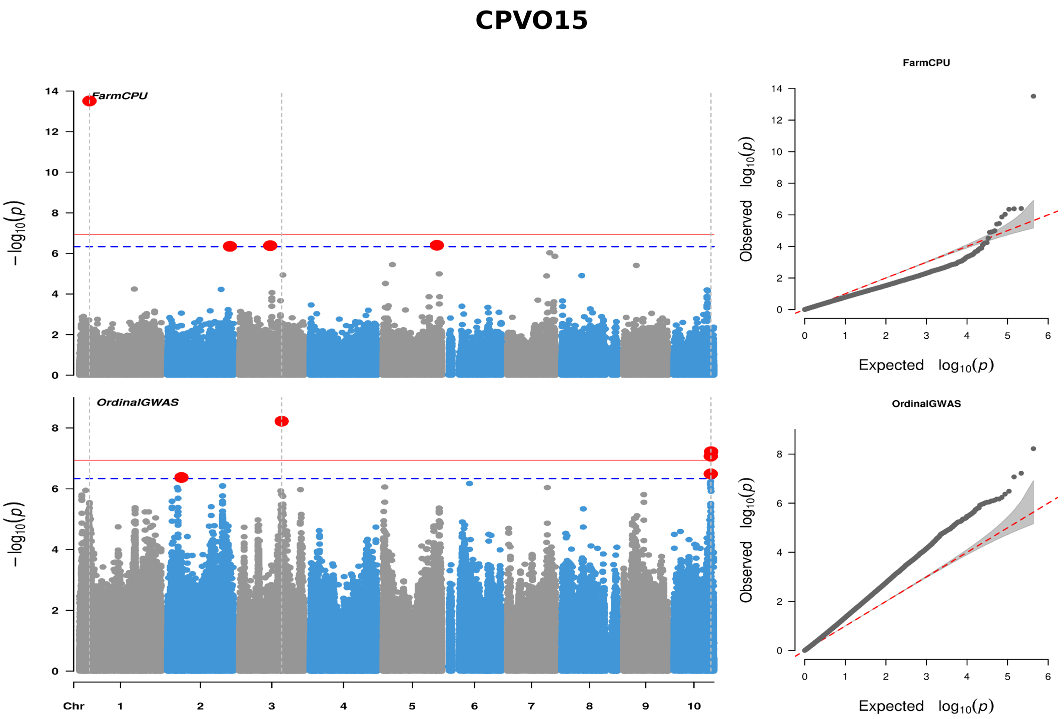
CPVO15
FarmCPU
OrdinalGWAS
−log10(p)
Chr
1 2 3 4 5 6 7 8 9 10
Observed log10(p)
Expected log10(p)

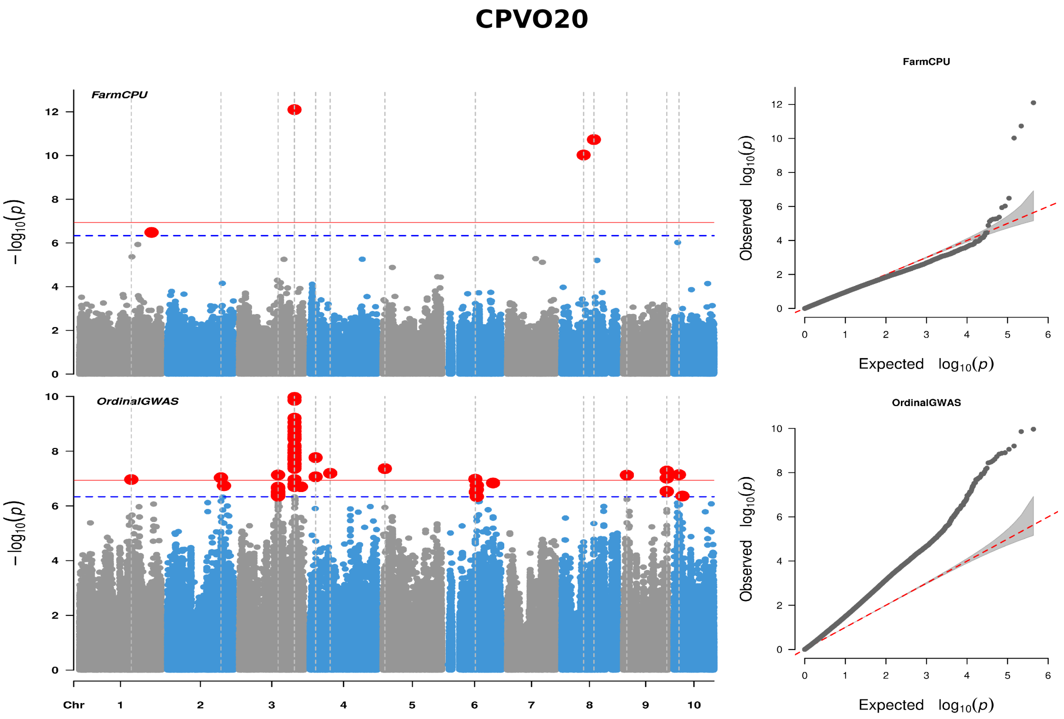
CPVO20
FarmCPU
OrdinalGWAS
−log10(p)
Chr
1 2 3 4 5 6 7 8 9 10
Observed log10(p)
Expected log10(p)

**CPVO26**

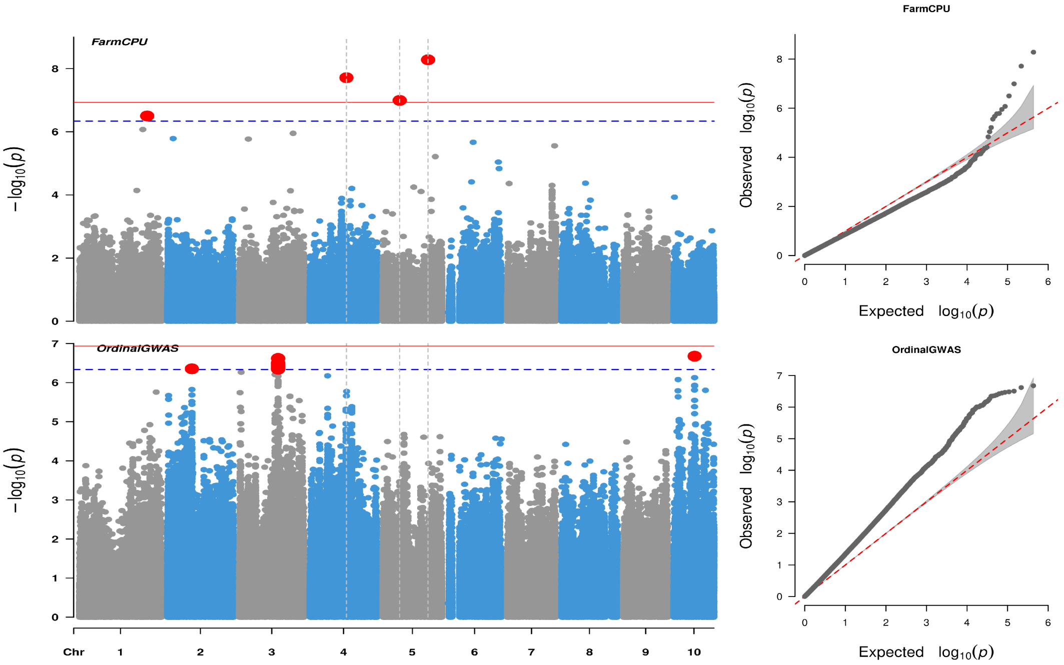


**CPVO29**

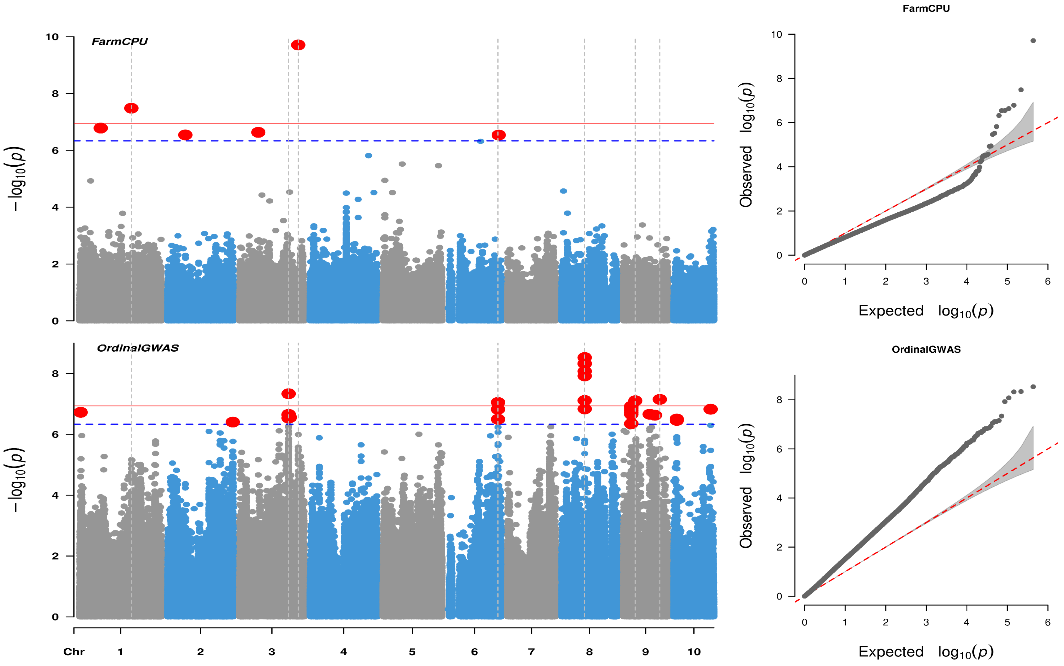

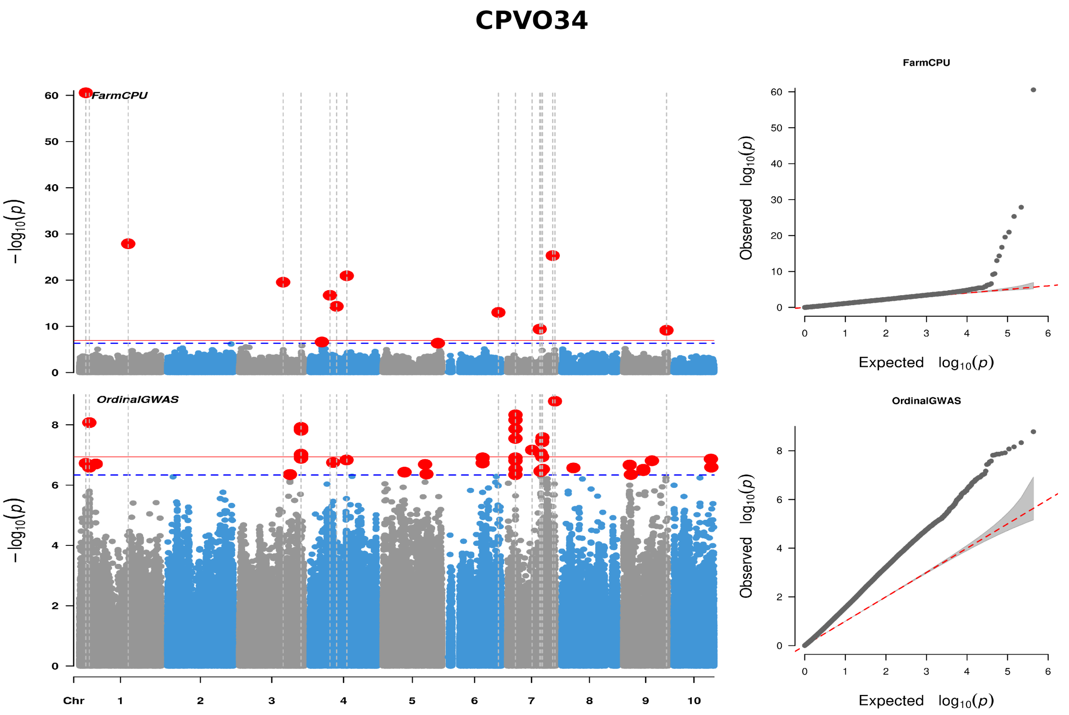


*Supplementary Figure 4.* Genome-wide association study (GWAS) results for DUS characters in the moderate-confidence subset: leaf: angle between blade and stem (CPVO4); tassel: anthocyanin coloration at base of glume (CPVO7); tassel: curvature of lateral branches (CPVO11); ear: anthocyanin coloration of silks (CPVO14); stem: anthocyanin coloration of brace roots (CPVO15); tassel: length of main axis above highest lateral branch (CPVO20); ear: length (CPVO26); ear: number of rows of grain (CPVO29); and ear: type of grain (CPVO34). Manhattan plots show genomic SNP positions versus $p$-values expressed as $-log_{10}(p)$ from the FarmCPU (top) and OrdinalGWAS analysis (bottom). The red solid line indicate the stringent Bonferroni significance threshold (FWER = 5% / $p = 1.16 \times 10^{-7}$), while the blue dashed line marks the suggestive threshold (FWER = 20% /$p = 4.62 \times 10^{-7}$). The vertical dashed lines depict SNPs which are significant (FWER = 5% / $p = 1.16 \times 10^{-7}$) with at least one of the two methods. The QQ plots compare observed and expected $p$-value distributions from the FarmCPU and OrdinalGWAS analysis.

### CPVO5

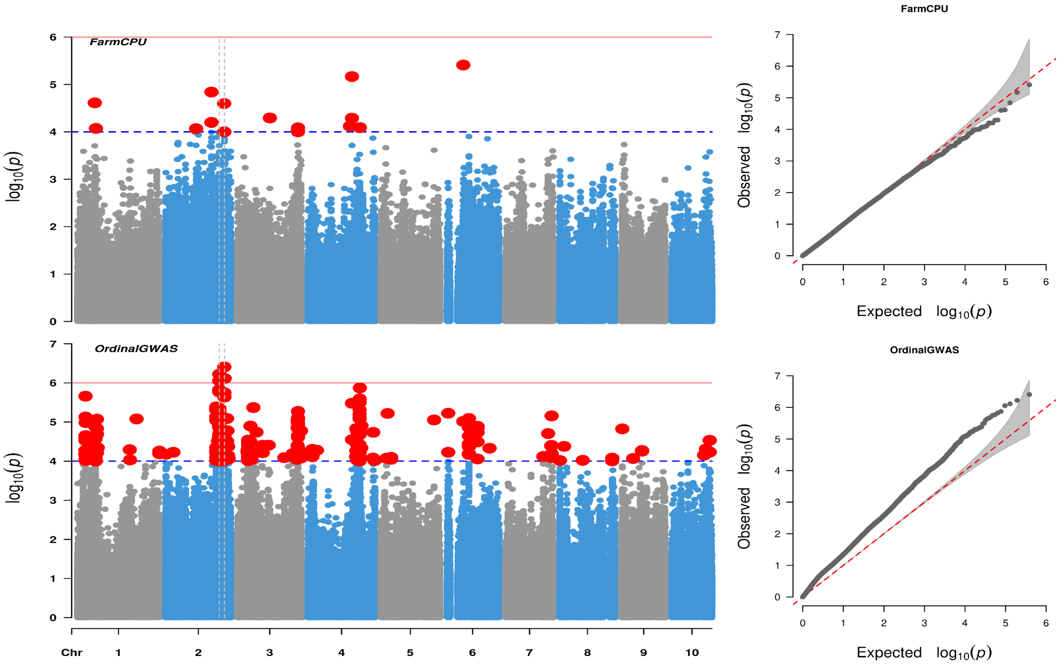


### CPVO22.2

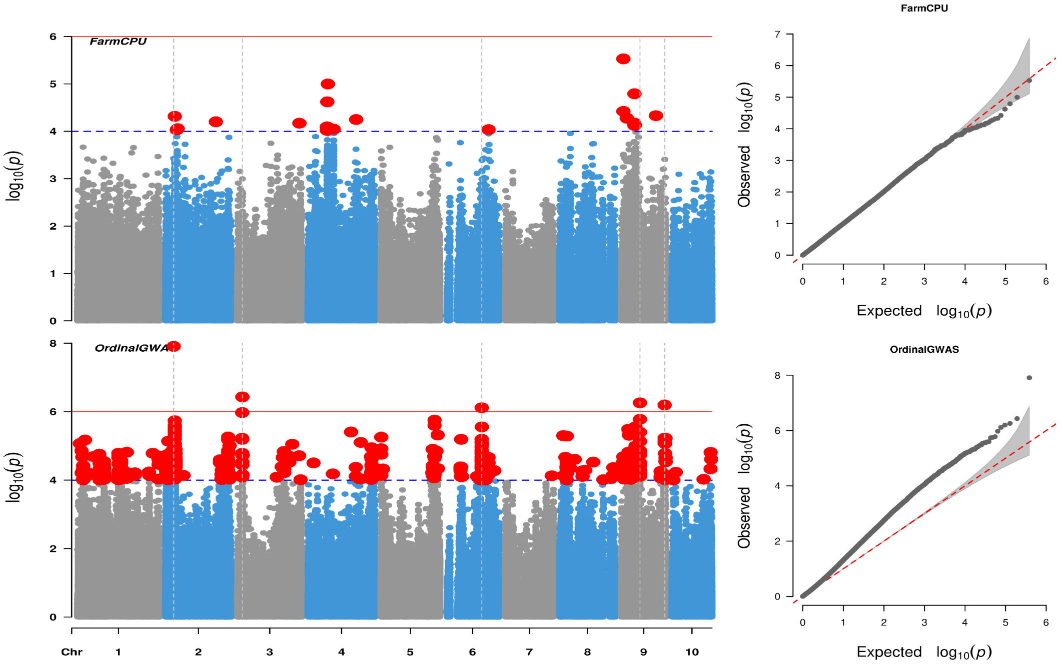

### CPVO23

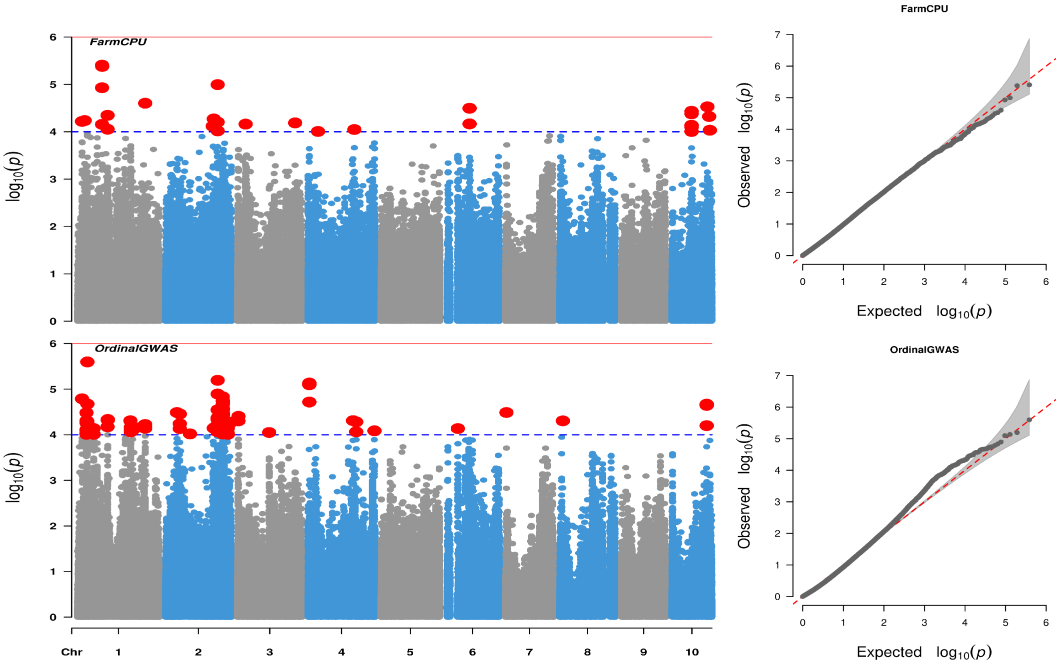


### CPVO27

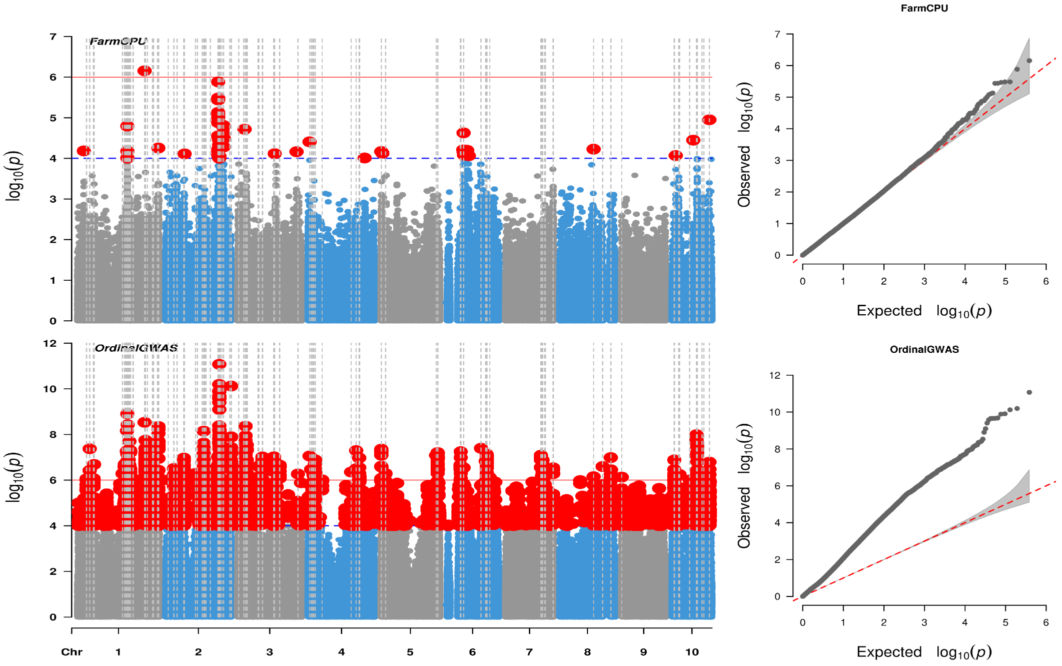

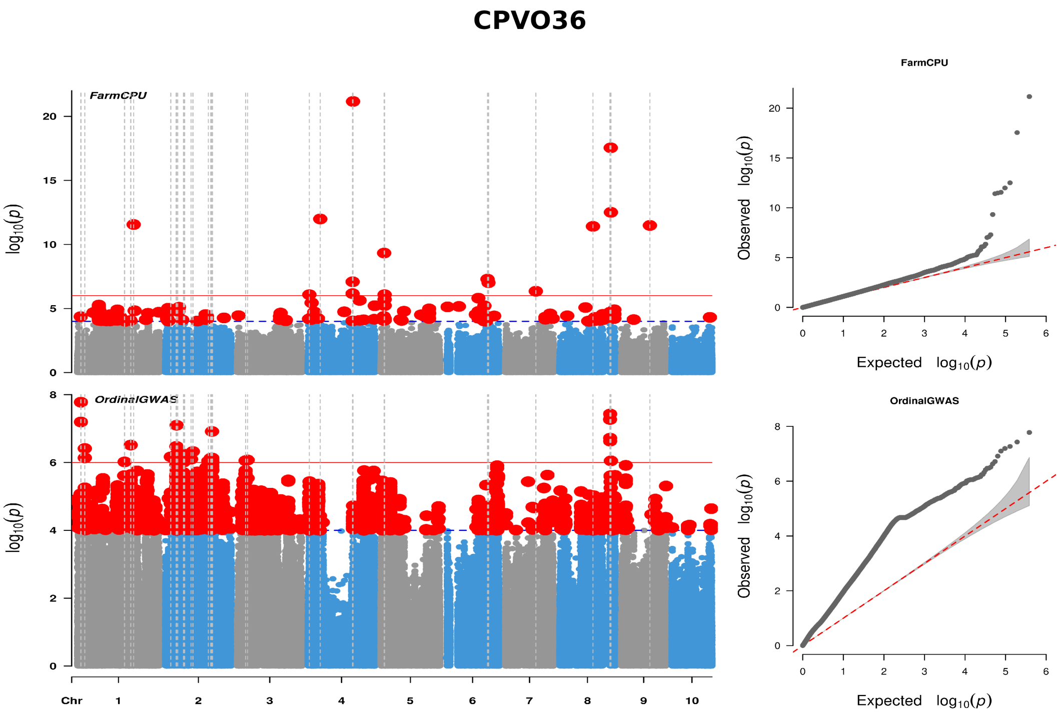


*Supplementary Figure 5.* Genome-wide association study (GWAS) results for DUS characters in the low-confidence subset: leaf: curvature of blade (CPVO5); plant: length (CPVO22.2) -evaluated only in hybrids and open-pollinated varieties, excluding those with sweet or pop grain types; plant: ratio of height of insertion of peduncle of upper ear to plant length (CPVO23); ear: diameter (middle) (CPVO27); and ear: colour of top of grain (CPVO36 ). In each Manhattan plots show genomic SNP positions versus $p$-values expressed as $-log_{10}(p)$ from the FarmCPU (top) and OrdinalGWAS analysis (bottom). The red solid line indicate the stringent Bonferroni significance threshold (FWER = 5% / $p = 1.30 \times 10^{-7}$), while the blue dashed line marks the suggestive threshold (FWER = 20% /$p = 5.21 \times 10^{-7}$). The vertical dashed lines depict SNPs which are significant (FWER = 5% / $p = 1.30 \times 10^{-7}$) with at least one of the two methods. The accompanying QQ plots compare observed and expected $p$-value distributions from the FarmCPU and OrdinalGWAS analysis.

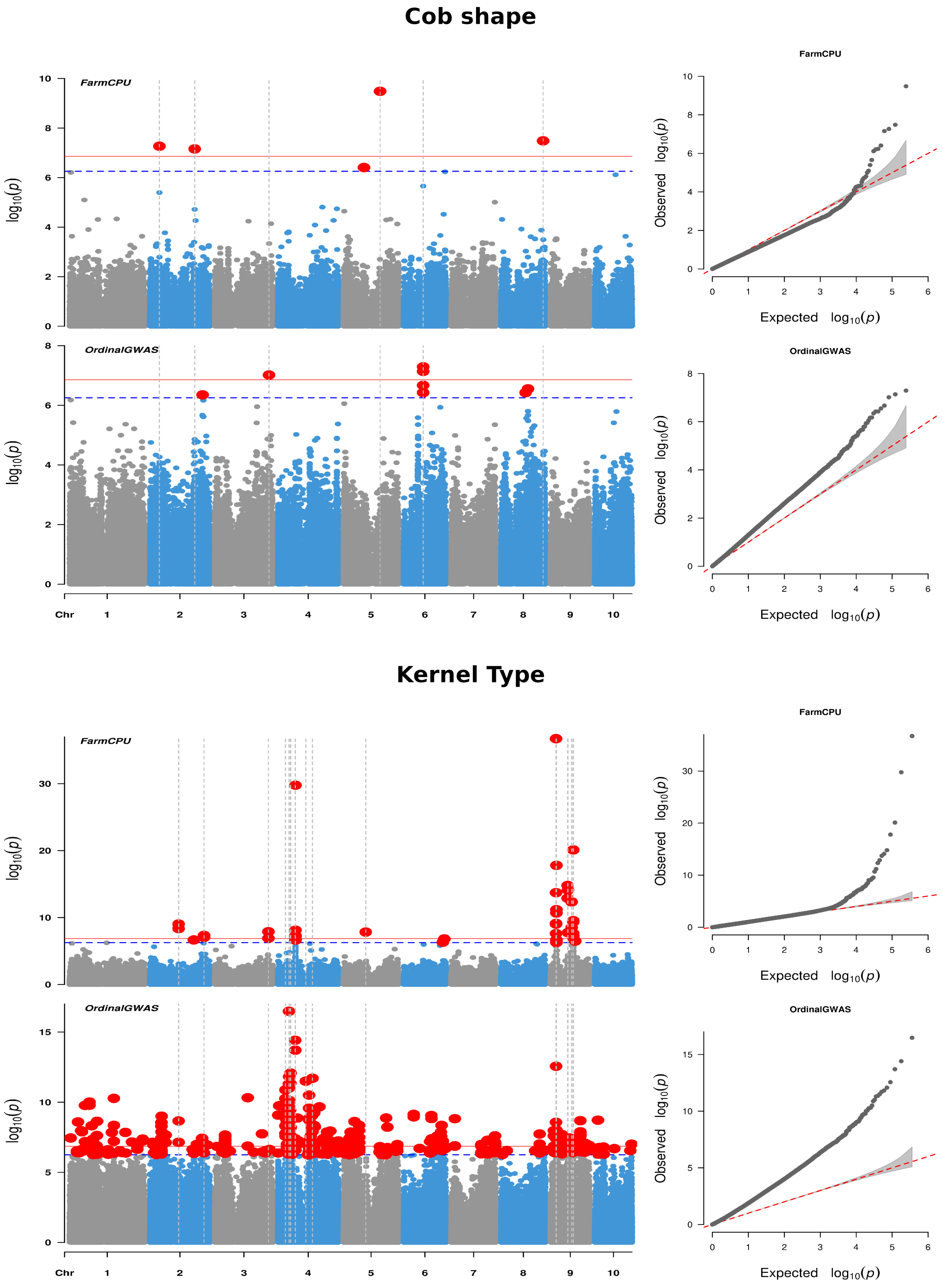


*Supplementary Figure 6.* Genome-wide association study (GWAS) analysis conducted using the USDA-NPGS dataset for "Cob shape" and "Kernel type". In each Manhattan plot show genomic SNP positions versus $p$-values expressed as $-log_{10}(p)$ from the FarmCPU (top) and OrdinalGWAS analysis (bottom). The red solid line indicate the stringent Bonferroni significance threshold (FWER = 5% / $p = 1.12 \times 10^{-7}$), while the blue dashed line marks the suggestive threshold (FWER = 20% /$p = 4.48 \times 10^{-4}$). The vertical dashed lines depict SNPs which are significant (FWER = 5% / $p = 1.16 \times 10^{-7}$) with at least one of the two methods.

The accompanying QQ plots compare observed and expected *p*-value distributions from the FarmCPU and OrdinalGWAS analysis.

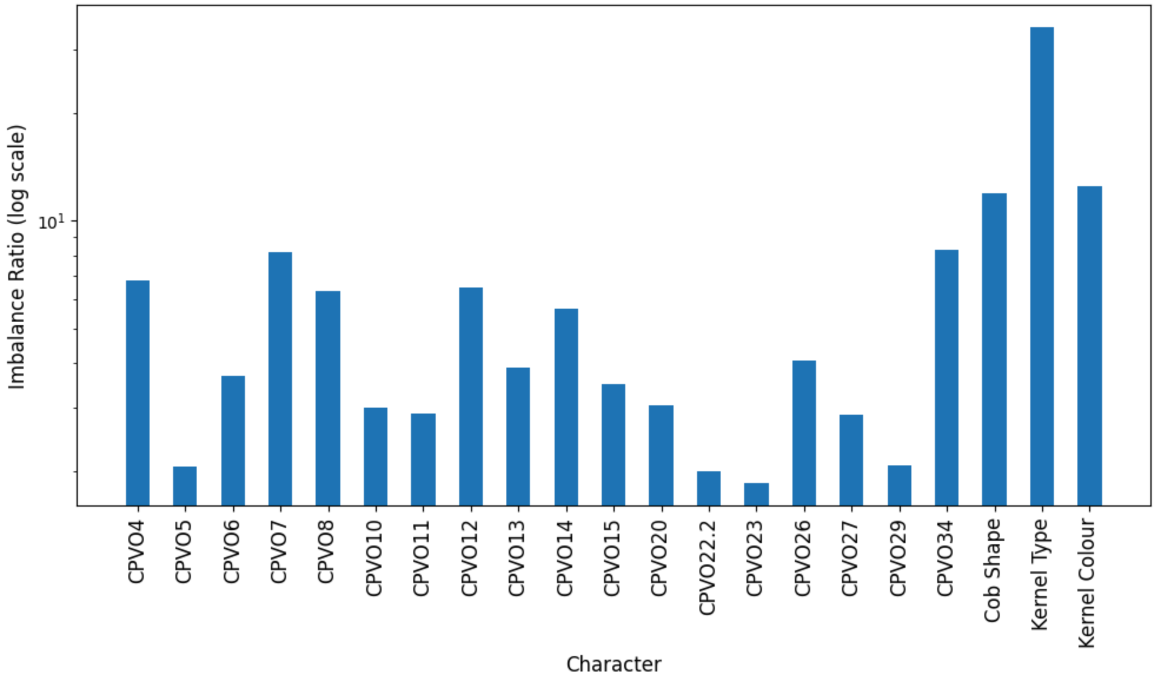


*Supplementary Figure 7.* Class imbalance ratios for the for 21 Distinctness, Uniformity, and Stability (DUS) characters displayed on a logarithmic scale. Character IDs correspond to 19 DUS characters from the European dataset as per CPVO testing protocol identifiers, along with three additional characters from the USDA-NPGS dataset: Cob Shape (CS), Kernel Colour (KC), and Kernel Type (KT). The class imbalance ratios here are calculated as ratio of the frequency of the majority class to the frequency of the minority class. A higher value indicates a greater disparity in category representation.
